# Preliminary report on SARS-CoV-2 Spike mutation T478K

**DOI:** 10.1101/2021.03.28.437369

**Authors:** Simone Di Giacomo, Daniele Mercatelli, Amir Rakhimov, Federico M. Giorgi

**Affiliations:** Department of Pharmacy and Biotechnology, University of Bologna, Via Selmi 3, 40138, Bologna, Italy

**Keywords:** SARS-CoV-2, COVID-19, Genomic Surveillance, Spike, T478K, S:T478K, Spike:T478K

## Abstract

Several SARS-CoV-2 variants have emerged, posing a renewed threat to COVID-19 containment and to vaccine and drug efficacy. In this study, we analyzed more than 1,000,000 SARS-CoV-2 genomic sequences deposited up to April 27, 2021 on the GISAID public repository, and identified a novel T478K mutation located on the SARS-CoV-2 Spike protein. The mutation is structurally located in the region of interaction with human receptor ACE2 and was detected in 11,435 distinct cases. We show that T478K has appeared and risen in frequency since January 2021, predominantly in Mexico and USA, but we could also detect it in several European countries.

## Introduction

Severe acute respiratory syndrome coronavirus 2 (SARS-CoV-2), the etiological cause of Coronavirus Disease 19 (COVID-19) is responsible for the most severe pandemic outbreak of the current century [1]. Naturally, it is the object of unprecedented scientific scrutiny, with more than one million SARS-CoV-2 genomic sequences having been generated and publicly shared since December 2019. This avalanche of data was made possible thanks to the efforts of thousands of contributing laboratories across the World, and collected by the GISAID initiative database [2]. This currently allows to run nearly real-time operations of genomic surveillance, by scrutinizing the evolution of the virus temporally and geographically [3]. In the first 17 months since the appearance of SARS-CoV-2, genomic surveillance has proven itself fundamental in tracking viral outbreaks [4] and in identifying potential new variants of clinical concern. One of these is the variant B.1.1.7 [5], characterized by 18 mutations over the reference genomic sequence (NCBI entry NC_045512.2, most notably a mutation A23063T, causing an aminoacidic change N501Y in the viral Spike protein interaction domain with human receptor Angiotensin Converting Enzyme 2 (ACE2) [6]. The interaction with ACE2, a surface protein expressed in human respiratory epithelial cells, is one of the key mechanisms for viral entry in the host, and it is a molecular mechanism directly connected with host specificity, early transmissibility [7] and higher viral infectivity [8].

N501Y is only one of the 9 Spike mutations of variant B.1.1.7, also characterized by mutations in polyprotein ORF1a, proteins ORF8 and Nucleocapsid (N) [9]. Another mutation in the Spike protein, D614G, has arisen in early 2020 and is currently present in more than 90% of all circulating SARS-CoV-2s; this mutation is not located in the interaction domain with ACE2, but it has been associated with increased entry efficiency with human host cells [10]. The US Center for Disease Control and Prevention (CDC) defined “variants of concern” all those mutations and lineages which have been associated with an increase in transmissibility and virulence, a decrease in the effectiveness of social and public health measures targeting the virus, and in general all mutations potentially affecting COVID-19 epidemiology [11]. Virtually all variants of concern contain mutations in the SARS-CoV-2 Spike protein, such as variant B.1.351 (Spike mutations K417N, E484K, N501Y, D614G and A701V) and variants B.1.427/B.1.429 (Spike mutations S13I, W152C, L452R and D614G), and many of these reside on the receptor binding domain (RBD), a region located between residues 350-550 of Spike and directly binding to the human ACE2 [12,13]. Recombinant vaccines comprising the Spike RBD induced potent antibody response in immunized animal models such as mouse, rabbits and primates [14]. On the other hand, mutations in the RBD can improve viral affinity to ACE2 and the evasion from neutralization antibodies [8,15–17]. A recently published clinical study [18] has shown also a decreased vaccine efficacy against the lineage B.1.351 (carrying Spike mutations E484K and N501Y), testifying the need to track and monitor all SARS-CoV-2 mutations, with a particular accent on those affecting the Spike RBD.

In this short Communication, we will show a report on a novel SARS-CoV-2 Spike mutation, T478K, which is also located at the interface of the Spike/ACE2 interaction, and it is worryingly rising in prevalence among SARS-CoV-2 sequences collected since the beginning of 2021.

## Materials and Methods

We downloaded all publicly available SARS-CoV-2 genomic sequences from the GISAID database on April 27, 2021. This yielded 1,180,571 samples, annotated with features such as collection date, region of origin, age, and sex of the infected patient. Only viruses collected from human hosts were kept for further processing, discarding e.g., environmental samples or viruses obtained from other mammals. We compared all these sequences with the SARS-CoV-2 Wuhan genome NC_045512.2, using a gene annotation file in GFF3 format available as Supplementary file 1. This provided 27,388,937 mutations when compared with the reference. These nucleotide mutations were then converted in corresponding cumulative effects on protein sequence using the Coronapp pipeline [19]. The 3D rendering of the location of S:T478K in the SARS-CoV-2 Spike / Human ACE2 complex was based on the crystal structure from [20], deposited in the Protein Data Bank [21] entry 6VW1. Comparison between wild-type and mutant Spike was performed using the Pymol suite [22] with the Adaptive Poisson–Boltzmann Solver (APBS) plugin [23]. All statistical analysis, algorithms and plotting were implemented with the R software [24].

## Results

In total, we could detect the Spike:T478K (S:T478K) mutation in 11,435 distinct patients as of April 27, 2021, more than twice the number observed one month before, on March 26, 2021 (4,214). The majority of these mutations (20,205 samples, Figure 1A and Supplementary Figure 1) are associated to PANGOLIN lineage B.1.1.519; S:T478K is present in 97.0% of B.1.1.519 cases. The remaining S:T478K events are distributed in small numbers (N<250) in other lineages phylogenetically not derived from B.1.1.519, supporting the hypothesis that this mutation has arisen more than once in distinct events. S:T478K is also present in 68 out of 85 (80%) reported samples from the B.1.214.3; however, the low total number of cases for this lineage do not make it a variant of concern yet.

**Figure 1.**
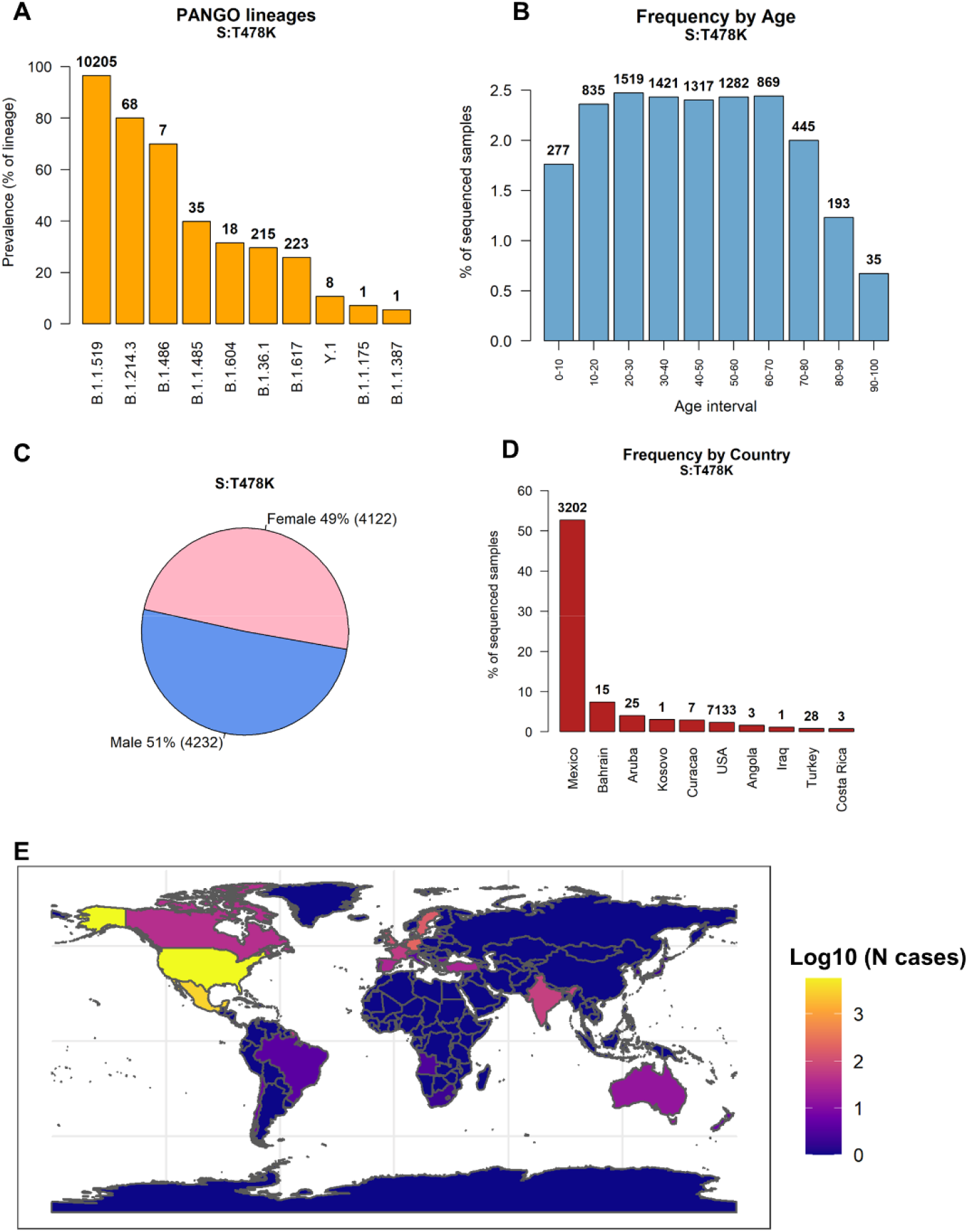
**A:** Prevalence of Spike mutation T478K in PANGO lineages. The top 10 lineages are reported, sorted by number of S:T478K samples over total number of lineage samples. The discrete number of S:T478K is reported on top of each bar. Most S:T478K-carrying samples are classified in the B.1.1.519 lineage. **B:** Frequency of sequenced SARS-CoV-2 genomes carrying the S:T478K mutation, divided into 10-years age ranges. The total number of S:T478K patients for the specified age range is reported on top of the bars. **C**: Pie chart showing the distribution of S:T478K by patient for which sex is reported. **D**: Number of S:T478K samples over total samples sequenced from each country. The 10 countries with higher frequency (in percentage) are shown. Discrete numbers of S:T478K are reported on top of each bar. **E**: Geographic global projection of S:T478K cases detected in each country. The color scale indicates number of SARS-CoV-2 genomes carrying the S:T478K mutation, in logarithm-10 scale.

S:T478K does not seem to be significantly associated with patient age (one way ANOVA test p>0.1, Figure 1B and Supplementary Figure 2), nor with patient sex (Figure 1C). The geographic distribution of S:T478K (Figure 1D and Supplementary Figure 3) shows a noticeable prevalence in Mexico, where it constitutes 52.8% (3,202 distinct cases) of all sequenced SARS-CoV-2 genomes. We could detect S:T478K mutations in 7,133 samples from the United States of America, totaling 2.7% of all genomes generated in the country. The S:T478K is therefore primarily present in North America, constituting more than 50% of all the sequences generated in Mexico (Figure 1D and Supplementary Figure 4). S:t478k has been detected also in European countries such as Germany, Sweden and Switzerland (Figure 1E and Supplementary Table 1).

One of the reasons of concern about S:T478K is that it is rapidly growing over time, both in number of detected samples (Figure 2A) and in prevalence, calculated as number of cases over total number of sequenced genomes (Figure 2B). We detected this grow starting at the beginning of 2021, and S:T478K is, at the time of writing (April 27, 2021) characterizing more than 2.0% of all sequenced SARS-CoV-2. As a comparison, we show the growth observed for Spike mutations S:N501Y, which rose in November 2020 (Figure 2C), and S:D614G, which exponentially grew in frequency starting February 2020 (Figure 2D).

**Figure 2.**
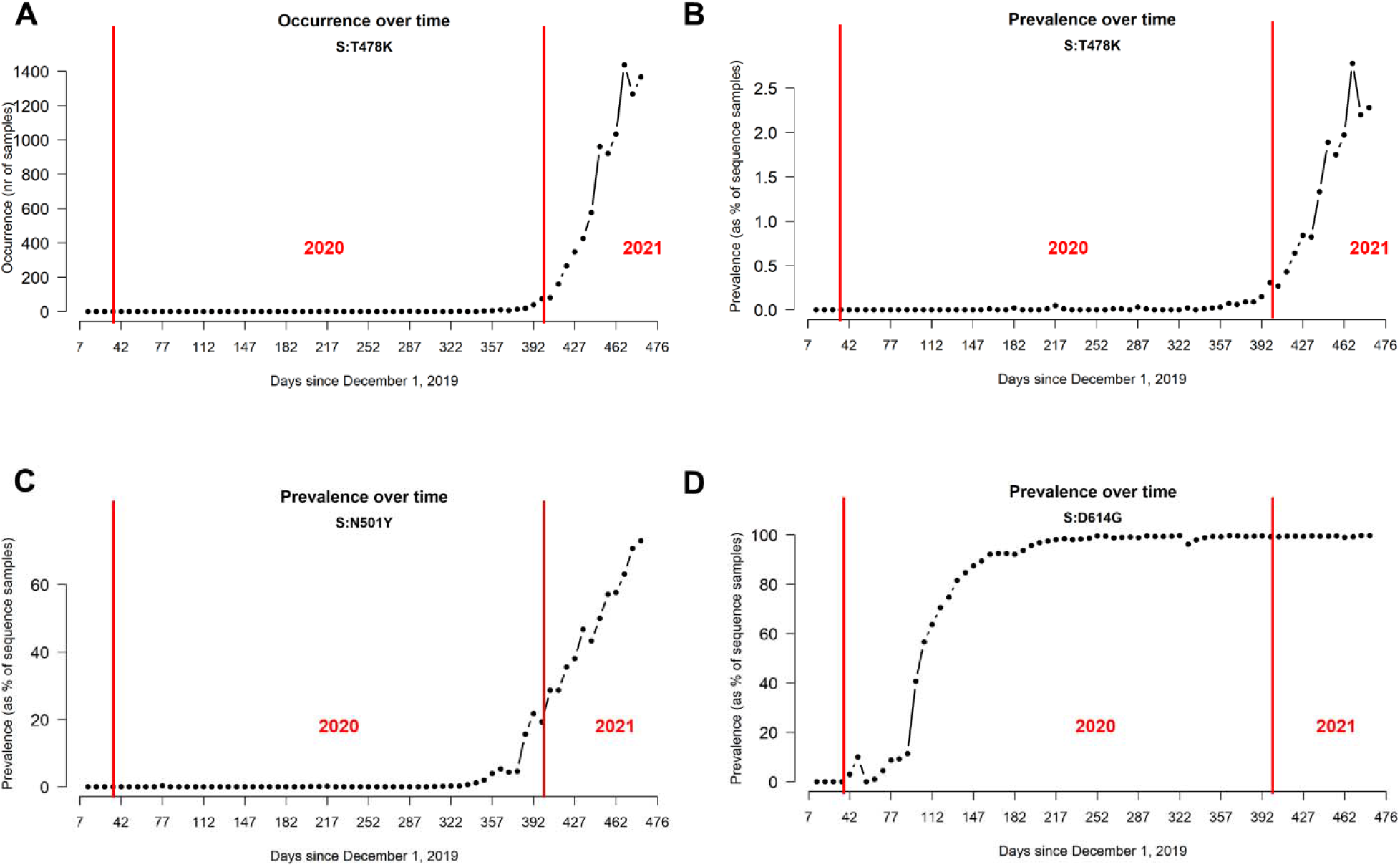
**A**: Number of sequenced SARS-CoV-2 genomes carrying S:T478K mutation over time, measured weekly. **B**: Prevalence over time of S:T478K in the SARS-CoV-2 population, measured as the number of S:T478K genomes over the total number of sequenced genomes. **C**: Prevalence over time of S:N501Y in the SARS-CoV-2 population. **D**: Prevalence over time of S:D614G in the SARS-CoV-2 population.

The location of S:T478K is within the interaction domain with the human receptor ACE2, roughly encompassing amino acids 350 to 550 of the SARS-CoV-2 Spike protein. In particular, the position of S:T478K is on the interface with ACE2, as shown by crystal structures of the complex (Figure 3A). The amino acid change from the polar but uncharged threonine (T) to a basic, charged lysine (K) is predicted to increase the electrostatic potential of Spike to a more positive surface, in a region directly contacting ACE2 (Figure 3B). Also, the larger side chain of lysine is predicted to increase the steric hindrance of the mutant, possibly further affecting the Spike/ACE2 interaction (Figure 3C).

**Figure 3.**
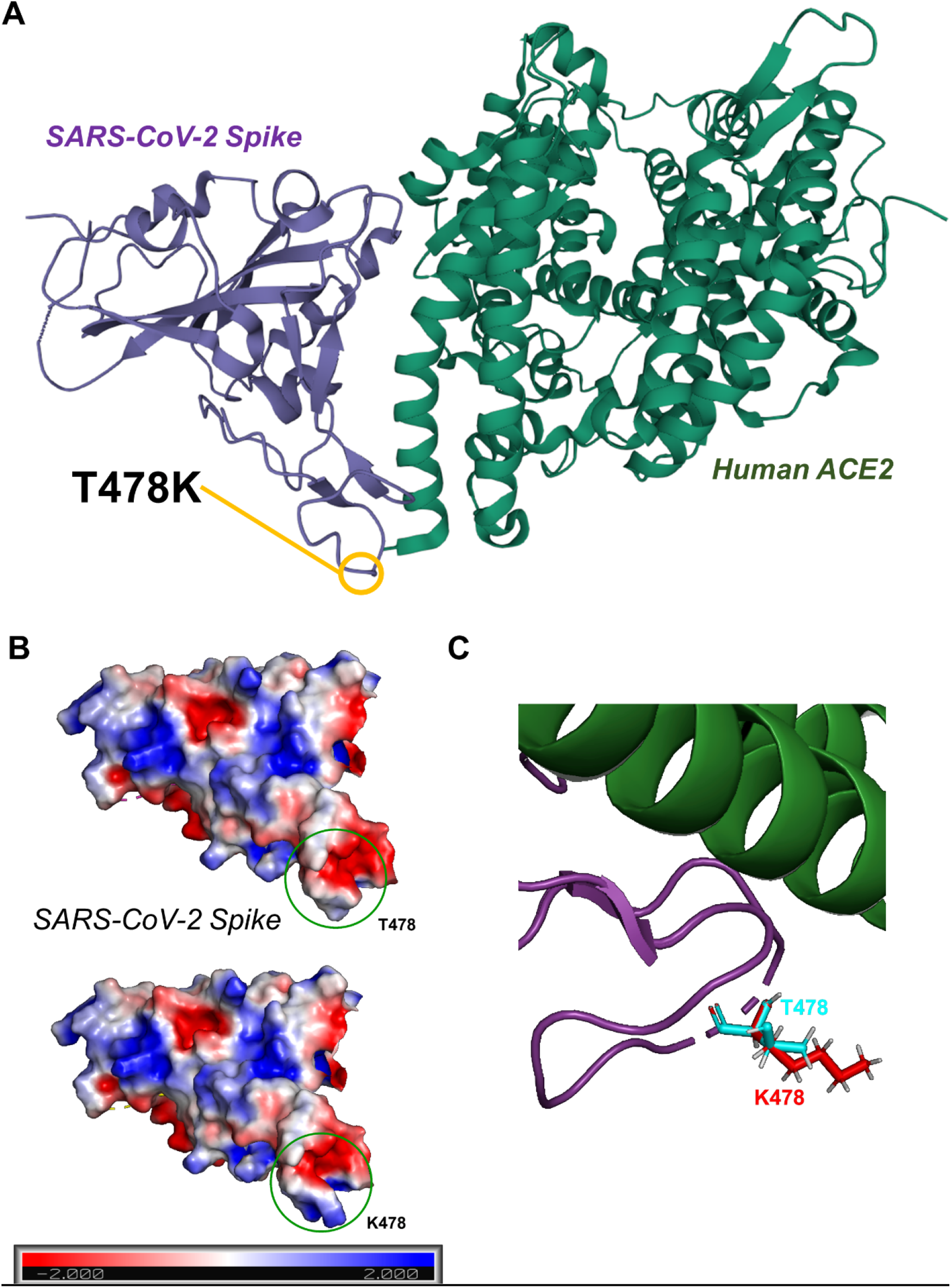
**A**: 3D representation of the SARS-CoV-2 Spike / Human ACE2 interacting complex, derived from the crystal structure from [20]. **B**: Representation of the SARS-CoV-2 Spike electrostatic potential calculated with the Adaptive Poisson-Boltzmann Solver (APBS) program [23] implemented in PyMOL [22]. Molecular surface was colored according to the molecular electrostatic potential [ranging from −2.0 (red) to 2.0 (blue)] in T478 (reference, above) and in K478 (more recent mutation, below). **C**: 3D detail of structural superposition of WT SARS-CoV-2 RBD and S:T478K. T478 side chains are colored in cyan, while K478 side chains are colored in red.

S:T478K is frequently co-occurring with three other Spike mutations located outside the canonical ACE2 interaction region. One is D614G (99.83% co-occurrence), one of the founding events of SARS-CoV-2 lineage B.*, currently the most diffused Worldwide (Table 1). The other two are P681H and T732A, with 93.8% and 88.7% co-occurrence with S:T478K, respectively (Table 1). We could detect S:T478K in co-presence with other Spike mutations as well, but currently all at much lower frequencies (<4%). The Spike S:T478K mutation is frequently co-existing also with mutations in other proteins, such as the diffused two-aa Nucleocapsid mutation N:RG203KR, and mutations in Non-Structural Proteins (NSPs) derived from the polyprotein encoded ORF1 (Open Reading Frame), which include for example the viral RNA-dependent RNA polymerase NSP12 (Table 2).

**Table 1.** Ten SARS-CoV-2 Spike mutations most frequently co-occurring with S:T478K. We report the number of genomes where both mutations are present, and the percentage over total number of samples where S:T478K is reported.

|  | Number of<br>cases | Percentage |
| --- | --- | --- |
| S:D614G | 11428 | 99.83 |
| S:P681H | 10739 | 93.81 |
| S:T732A | 10157 | 88.73 |
| S:L5F | 439 | 3.84 |
| S:L452R | 241 | 2.11 |
| S:P681R | 224 | 1.96 |
| S:D950N | 222 | 1.94 |
| S:T19R | 215 | 1.88 |
| S:E156 | 174 | 1.52 |
| S:E154A | 149 | 1.3 |

**Table 2.** Ten SARS-CoV-2 Non-Spike mutations most frequently co-occurring with S:T478K. We report the number of genomes where both mutations are present, and the percentage over total number of samples where S:7478K is reported. NSP: Non-Structural Protein; N: Nucleocapsid protein; ORF: Open Reading Frame.

|  | Number of<br>cases | Percentage |
| --- | --- | --- |
| NSP12b:P314L | 11310 | 98.8 |
| NSP4:T492I | 10716 | 93.61 |
| NSP6:I49V | 10692 | 93.4 |
| NSP3:P141S | 10687 | 93.36 |
| N:RG203KR | 10622 | 92.79 |
| NSP9:T35I | 10150 | 88.67 |
| ORF8:L4P | 1617 | 14.13 |
| N:Q418H | 1052 | 9.19 |
| NSP2:T44I | 1044 | 9.12 |
| ORF8:A65V | 616 | 5.38 |

## Discussion

In this short communication, we report the distribution of the Spike mutation S:T478K and its recent growth in prevalence in the SARS-CoV-2 population. While there is currently no report of association of this variant with clinical features, S:T478K’s rapid growth may indicate an increased adaption of SARS-CoV-2 variants carrying it, particularly lineage B.1.1.519. The distribution of this mutation, which emerged from the B.1 lineage carrying S:D614G, but is independent from the S:N501Y mutation, is higher in North America [25], but we could detect it also in several European countries. T478K has been detected in other phylogenetically non-derived lineages from B.1.1.519, supporting the hypothesis that this mutation arose more than once in distinct events. Since the highest abundance of this mutation seems to be in Mexico and USA, this may allow to hypothesize a founder effect in which a chance founder event was followed by natural selection progression, since the frequency of the mutation has, slowly but steadily, increased in the first months of 2021.

The location of S:T478K in the interaction complex with human ACE2 may affect the affinity with human cells and therefore influence viral infectivity. An **in silico** molecular dynamics study on the protein structure of Spike has predicted that the T478K mutation, substituting a non-charged amino acid (Threonine) with a positive one (Lysine) may significantly alter the electrostatic surface of the protein (Figure 3), and therefore the interaction with ACE2, drugs or antibodies [26], and that the effect can be increased if combined by other co-occurring Spike mutations (see Table 1). Another experiment showed that T478K and T478R mutants were enriched when SARS-CoV-2 viral cultures were tested against weak neutralizing antibodies [27], highlighting, at least **in vitro**, a possible genetic route the virus can follow to escape immune recognition. Everything considered, we believe that the continued genetical and clinical monitoring of S:T478K and other Spike mutations is of paramount importance to better understand COVID-19 and be able to better counteract its future developments.

## Supplementary Material

Supplementary file 1: the SARS-CoV-2 genome annotation coordinates, in GFF3 format, used in this study, and based on NCBI reference genome sequence NC_045512.2.

Supplementary file 2: text file listing the original contributors of sequences analyzed in this study, and deposited on the GISAID database.

Supplementary figure 1: nr of samples carrying the S:T478K mutation, divided by lineages. Supplementary figure 2: nr of samples carrying the S:T478K mutation, divided by age ranges.

Supplementary figure 3: 10 countries with the highest nr of samples carrying the S:T478K mutation.

Supplementary figure 4: world projection showing the percentage frequency of S:T478K over the total number of sequenced samples.

Supplementary table 1: Excel table showing the number of S:T478K in European countries, both absolute and as a percentage of the sequenced samples.

## Author Contributions

Conceptualization, SDG and FMG. Funding acquisition, FMG. Writing – original draft preparation, FMG. Writing – review and editing, DM and SDG. Methodology, AR and FMG. Validation, DM, SDG and FMG. Software, AR and FMG. All authors contributed to the study and approved the final version of the manuscript.

## Data Availability Statement

All data supporting this study is available on the GISAID portal https://www.gisaid.org/

## Acknowledgments

We are very grateful to the GISAID Initiative and all its data contributors, i.e. the Authors from the originating laboratories responsible for obtaining the specimens and the Submitting laboratories where genetic sequence data were generated and shared via the GISAID Initiative, on which this research is based.

