## Supplementary material for "Preliminary report on SARS-CoV-2 Spike mutation T478K": SuppFile1_NC_045512.2_annot.gff3.docx

##sequence-region NC_045512.2 1 29903

##species https://www.ncbi.nlm.nih.gov/Taxonomy/Browser/wwwtax.cgi?id=2697049

NC_045512.2	Giorgi	CDS	266	805	.	+	.	NSP1	Leader protein

NC_045512.2	Giorgi	CDS	806	2719	.	+	.	NSP2	Non-Structural protein 2

NC_045512.2	Giorgi	CDS	2720	8554	.	+	.	NSP3	Predicted phosphoesterase, papain-like proteinase

NC_045512.2	Giorgi	CDS	8555	10054	.	+	.	NSP4	Transmembrane protein

NC_045512.2	Giorgi	CDS	10055	10972	.	+	.	NSP5	3C-like proteinase

NC_045512.2	Giorgi	CDS	10973	11842	.	+	.	NSP6	Transmembrane protein

NC_045512.2	Giorgi	CDS	11843	12091	.	+	.	NSP7	Non-Structural Protein 7

NC_045512.2	Giorgi	CDS	12092	12685	.	+	.	NSP8	Non-Structural Protein 8

NC_045512.2	Giorgi	CDS	12686	13024	.	+	.	NSP9	ssRNA-binding protein

NC_045512.2	Giorgi	CDS	13025	13441	.	+	.	NSP10	Growth-factor-like protein

NC_045512.2	Giorgi	CDS	13442	13468	.	+	.	NSP12a	RNA-dependent RNA polymerase, pre-ribosomal frameshift

NC_045512.2	Giorgi	CDS	13468	16236	.	+	.	NSP12b	RNA-dependent RNA polymerase, post-ribosomal frameshift

NC_045512.2	Giorgi	CDS	16237	18039	.	+	.	NSP13	Helicase

NC_045512.2	Giorgi	CDS	18040	19620	.	+	.	NSP14	3'-to-5' exonuclease

NC_045512.2	Giorgi	CDS	19621	20658	.	+	.	NSP15	endoRNAse

NC_045512.2	Giorgi	CDS	20659	21552	.	+	.	NSP16	2'-O-ribose methyltransferase

NC_045512.2	Giorgi	CDS	21563	25384	.	+	.	S	Spike

NC_045512.2	Giorgi	CDS	25393	26220	.	+	.	ORF3a	ORF3a protein

NC_045512.2	Giorgi	CDS	26245	26472	.	+	.	E	Envelope

NC_045512.2	Giorgi	CDS	26523	27191	.	+	.	M	Membrane

NC_045512.2	Giorgi	CDS	27202	27387	.	+	.	ORF6	ORF6 protein

NC_045512.2	Giorgi	CDS	27394	27759	.	+	.	ORF7a	ORF7a protein

NC_045512.2	Giorgi	CDS	27756	27887	.	+	.	ORF7b	ORF7b protein

NC_045512.2	Giorgi	CDS	27894	28259	.	+	.	ORF8	ORF8 protein

NC_045512.2	Giorgi	CDS	28274	29533	.	+	.	N	Nucleocapsid protein

NC_045512.2	Giorgi	CDS	29558	29674	.	+	.	ORF10	ORF10 protein
