## Supplementary material for "Preliminary report on SARS-CoV-2 Spike mutation T478K": SuppFile2_sources.docx

"1. Genome Research Center for Health (CRGS) / 2. Laboratory of Molecular Medicine and Genomics(LMMGe) / 3. Center for Research in Pure and Applied Mathematics (CRMPA)"

"School of Life Sciences and Technology & School of Pharmacy-Institut Teknologi Bandung; Molecular Genetics Laboratory-Faculty of Medicine-Universitas Padjadjaran; Laboratorium Kesehatan Provinsi Jawa Barat"

"Stefan cel Mare" University Metagenomics Lab

"Stefan cel Mare" University Metagenomics Laboratory

"Stefan S. Nicolau"Institute of Virology

"Swiss Tropical and Public Health Institute"

"Unité Mixte Internationale TransVIHMI (UMI 233 IRD – U1175 INSERM - Université de Montpellier) IRD (Institut de recherche pour le développement)"

[Romania, Bucharest] National Institute for Infectious Diseases “Prof. Dr. Matei Balș”

Evandro Chagas Institute

0. Laboratory of Recombinant Vaccines, Intercollegiate Faculty of Biotechnology University of Gdansk and Medical University of Gdansk, 2. ViroGenetics - BSL3 Laboratory of Virology, Malopolska Centre of Biotechnology, Jagiellonian University

1-Clinical and Experimental Pharmacology Lab, LR16SP02, National Center of Pharmacovigilance, University of Tunis El Manar, Tunis, Tunisia. 2-Neurodegenerative diseases and psychiatric troubles, LR18SP03, Razi Hospital, University of Tunis El Manar, Tunis, Tunisia. 3- Ministry of Health, National Observatory of New and Emerging Diseases, 1006, Tunis, Tunisia

1-Laboratory of Microbiology, National Reference Lab, Charles Nicolle Hospital; 2-University of Tunis ElManar, Faculty of Medicine of Tunis, LR99ES09, Tunis, Tunisia

1) Dept. Infectious, Tropical Diseases & Microbiology, IRCCS Sacro Cuore Don Calabria Hospital; 2) Centro Piattaforme Tecnologiche, University of Verona; 3) Dept. Neurosciences, Biomedicine and Movement Sciences, University of Verona.

1. Academic Center for Pathomorphological and Genetic-Molecular Diagnostics ltd, Bialystok, Poland 2. National Institute of Public Health - National Institute of Hygiene, Warsaw, Poland

1. Department of Medical Sciences, Ministry of Public Health, Thailand 2. Thai Red Cross Emerging Infectious Diseases - Health Science Centre 3. Department of Disease Control, Ministry of Public Health, Thailand

1. Genome Research Center for Health (CRGS) / 2. Laboratory of Molecular Medicine and Genomics(LMMGe) / 3. Center for Research in Pure and Applied Mathematics (CRMPA)

1. Laboratory of Communicable Diseases (Estonia); 2. Eurofins Genomics Europe Sequencing GmbH

1. Laboratory of Recombinant Vaccines 2. Department of Biology and Medical Genetics 3. Laboratory of Clinical Genetics

1. Laboratory of Recombinant Vaccines, Intercollegiate Faculty of Biotechnology University of Gdansk and Medical University of Gdansk, 2. ViroGenetics - BSL3 Laboratory of Virology, Malopolska Centre of Biotechnology, Jagiellonian University

1. National Institute of Public Health - National Institute of Hygiene; 2. Eurofins Genomics Europe Sequencing GmbH

1. ViroGenetics - BSL3 Laboratory of Virology, Malopolska Centre of Biotechnology, Jagiellonian University; 2. Diagtron Laboratoria Lukasz Rabalski

1. ViroGenetics - BSL3 Laboratory of Virology, Malopolska Centre of Biotechnology, Jagiellonian University; 2. genXone SA, Research & Development Laboratory

1. ViroGenetics - BSL3 Laboratory of Virology, Malopolska Centre of Biotechnology, Jagiellonian University; 2. Human Genome Variation Research Group, Malopolska Centre of Biotechnology, Jagiellonian University;

1. ViroGenetics - BSL3 Laboratory of Virology, Malopolska Centre of Biotechnology, Jagiellonian University; 2. II Department of Internal Medicine, Faculty of Medicine, Jagiellonian University Medical College.

1. ViroGenetics - BSL3 Laboratory of Virology, Malopolska Centre of Biotechnology, Jagiellonian University; 2. II Department of Internal Medicine, Faculty of Medicine, Jagiellonian University Medical College; 3. Narodowy Instytut Zdrowia Publicznego – Panstwowy Zaklad Higieny (NIZP-PZH).

1. Virogenetics Laboratory of Virology, Malopolska Centre of Biotechnology, Jagiellonian University. 2. Intercollegiate Faculty of Biotechnology University of Gdansk and Medical University of Gdansk

10. Laboratory of Recombinant Vaccines, Intercollegiate Faculty of Biotechnology University of Gdansk and Medical University of Gdansk, 2. ViroGenetics - BSL3 Laboratory of Virology, Malopolska Centre of Biotechnology, Jagiellonian University

11. Laboratory of Recombinant Vaccines, Intercollegiate Faculty of Biotechnology University of Gdansk and Medical University of Gdansk, 2. ViroGenetics - BSL3 Laboratory of Virology, Malopolska Centre of Biotechnology, Jagiellonian University

12. Laboratory of Recombinant Vaccines, Intercollegiate Faculty of Biotechnology University of Gdansk and Medical University of Gdansk, 2. ViroGenetics - BSL3 Laboratory of Virology, Malopolska Centre of Biotechnology, Jagiellonian University

1270 Natividad Road Salinas, CA 93906

13. Laboratory of Recombinant Vaccines, Intercollegiate Faculty of Biotechnology University of Gdansk and Medical University of Gdansk, 2. ViroGenetics - BSL3 Laboratory of Virology, Malopolska Centre of Biotechnology, Jagiellonian University

14. Laboratory of Recombinant Vaccines, Intercollegiate Faculty of Biotechnology University of Gdansk and Medical University of Gdansk, 2. ViroGenetics - BSL3 Laboratory of Virology, Malopolska Centre of Biotechnology, Jagiellonian University

15. Laboratory of Recombinant Vaccines, Intercollegiate Faculty of Biotechnology University of Gdansk and Medical University of Gdansk, 2. ViroGenetics - BSL3 Laboratory of Virology, Malopolska Centre of Biotechnology, Jagiellonian University

16. Laboratory of Recombinant Vaccines, Intercollegiate Faculty of Biotechnology University of Gdansk and Medical University of Gdansk, 2. ViroGenetics - BSL3 Laboratory of Virology, Malopolska Centre of Biotechnology, Jagiellonian University

17. Laboratory of Recombinant Vaccines, Intercollegiate Faculty of Biotechnology University of Gdansk and Medical University of Gdansk, 2. ViroGenetics - BSL3 Laboratory of Virology, Malopolska Centre of Biotechnology, Jagiellonian University

18. Laboratory of Recombinant Vaccines, Intercollegiate Faculty of Biotechnology University of Gdansk and Medical University of Gdansk, 2. ViroGenetics - BSL3 Laboratory of Virology, Malopolska Centre of Biotechnology, Jagiellonian University

19-21, boulevard Jean Moulin, 13005 Marseille

19. Laboratory of Recombinant Vaccines, Intercollegiate Faculty of Biotechnology University of Gdansk and Medical University of Gdansk, 2. ViroGenetics - BSL3 Laboratory of Virology, Malopolska Centre of Biotechnology, Jagiellonian University

2. Laboratory of Recombinant Vaccines, Intercollegiate Faculty of Biotechnology University of Gdansk and Medical University of Gdansk, 2. ViroGenetics - BSL3 Laboratory of Virology, Malopolska Centre of Biotechnology, Jagiellonian University

20. Laboratory of Recombinant Vaccines, Intercollegiate Faculty of Biotechnology University of Gdansk and Medical University of Gdansk, 2. ViroGenetics - BSL3 Laboratory of Virology, Malopolska Centre of Biotechnology, Jagiellonian University

21. Laboratory of Recombinant Vaccines, Intercollegiate Faculty of Biotechnology University of Gdansk and Medical University of Gdansk, 2. ViroGenetics - BSL3 Laboratory of Virology, Malopolska Centre of Biotechnology, Jagiellonian University

22. Laboratory of Recombinant Vaccines, Intercollegiate Faculty of Biotechnology University of Gdansk and Medical University of Gdansk, 2. ViroGenetics - BSL3 Laboratory of Virology, Malopolska Centre of Biotechnology, Jagiellonian University

23. Laboratory of Recombinant Vaccines, Intercollegiate Faculty of Biotechnology University of Gdansk and Medical University of Gdansk, 2. ViroGenetics - BSL3 Laboratory of Virology, Malopolska Centre of Biotechnology, Jagiellonian University

24. Laboratory of Recombinant Vaccines, Intercollegiate Faculty of Biotechnology University of Gdansk and Medical University of Gdansk, 2. ViroGenetics - BSL3 Laboratory of Virology, Malopolska Centre of Biotechnology, Jagiellonian University

25. Laboratory of Recombinant Vaccines, Intercollegiate Faculty of Biotechnology University of Gdansk and Medical University of Gdansk, 2. ViroGenetics - BSL3 Laboratory of Virology, Malopolska Centre of Biotechnology, Jagiellonian University

26. Laboratory of Recombinant Vaccines, Intercollegiate Faculty of Biotechnology University of Gdansk and Medical University of Gdansk, 2. ViroGenetics - BSL3 Laboratory of Virology, Malopolska Centre of Biotechnology, Jagiellonian University

27. Laboratory of Recombinant Vaccines, Intercollegiate Faculty of Biotechnology University of Gdansk and Medical University of Gdansk, 2. ViroGenetics - BSL3 Laboratory of Virology, Malopolska Centre of Biotechnology, Jagiellonian University

28. Laboratory of Recombinant Vaccines, Intercollegiate Faculty of Biotechnology University of Gdansk and Medical University of Gdansk, 2. ViroGenetics - BSL3 Laboratory of Virology, Malopolska Centre of Biotechnology, Jagiellonian University

285 Mihai Bravu Ave, Bucharest, Romania

29. Laboratory of Recombinant Vaccines, Intercollegiate Faculty of Biotechnology University of Gdansk and Medical University of Gdansk, 2. ViroGenetics - BSL3 Laboratory of Virology, Malopolska Centre of Biotechnology, Jagiellonian University

3. Laboratory of Recombinant Vaccines, Intercollegiate Faculty of Biotechnology University of Gdansk and Medical University of Gdansk, 2. ViroGenetics - BSL3 Laboratory of Virology, Malopolska Centre of Biotechnology, Jagiellonian University

30. Laboratory of Recombinant Vaccines, Intercollegiate Faculty of Biotechnology University of Gdansk and Medical University of Gdansk, 2. ViroGenetics - BSL3 Laboratory of Virology, Malopolska Centre of Biotechnology, Jagiellonian University

31. Laboratory of Recombinant Vaccines, Intercollegiate Faculty of Biotechnology University of Gdansk and Medical University of Gdansk, 2. ViroGenetics - BSL3 Laboratory of Virology, Malopolska Centre of Biotechnology, Jagiellonian University

32. Laboratory of Recombinant Vaccines, Intercollegiate Faculty of Biotechnology University of Gdansk and Medical University of Gdansk, 2. ViroGenetics - BSL3 Laboratory of Virology, Malopolska Centre of Biotechnology, Jagiellonian University

33. Laboratory of Recombinant Vaccines, Intercollegiate Faculty of Biotechnology University of Gdansk and Medical University of Gdansk, 2. ViroGenetics - BSL3 Laboratory of Virology, Malopolska Centre of Biotechnology, Jagiellonian University

35. Laboratory of Recombinant Vaccines, Intercollegiate Faculty of Biotechnology University of Gdansk and Medical University of Gdansk, 2. ViroGenetics - BSL3 Laboratory of Virology, Malopolska Centre of Biotechnology, Jagiellonian University

37. Laboratory of Recombinant Vaccines, Intercollegiate Faculty of Biotechnology University of Gdansk and Medical University of Gdansk, 2. ViroGenetics - BSL3 Laboratory of Virology, Malopolska Centre of Biotechnology, Jagiellonian University

38. Laboratory of Recombinant Vaccines, Intercollegiate Faculty of Biotechnology University of Gdansk and Medical University of Gdansk, 2. ViroGenetics - BSL3 Laboratory of Virology, Malopolska Centre of Biotechnology, Jagiellonian University

39. Laboratory of Recombinant Vaccines, Intercollegiate Faculty of Biotechnology University of Gdansk and Medical University of Gdansk, 2. ViroGenetics - BSL3 Laboratory of Virology, Malopolska Centre of Biotechnology, Jagiellonian University

4. Laboratory of Recombinant Vaccines, Intercollegiate Faculty of Biotechnology University of Gdansk and Medical University of Gdansk, 2. ViroGenetics - BSL3 Laboratory of Virology, Malopolska Centre of Biotechnology, Jagiellonian University

40. Laboratory of Recombinant Vaccines, Intercollegiate Faculty of Biotechnology University of Gdansk and Medical University of Gdansk, 2. ViroGenetics - BSL3 Laboratory of Virology, Malopolska Centre of Biotechnology, Jagiellonian University

41. Laboratory of Recombinant Vaccines, Intercollegiate Faculty of Biotechnology University of Gdansk and Medical University of Gdansk, 2. ViroGenetics - BSL3 Laboratory of Virology, Malopolska Centre of Biotechnology, Jagiellonian University

42. Laboratory of Recombinant Vaccines, Intercollegiate Faculty of Biotechnology University of Gdansk and Medical University of Gdansk, 2. ViroGenetics - BSL3 Laboratory of Virology, Malopolska Centre of Biotechnology, Jagiellonian University

43. Laboratory of Recombinant Vaccines, Intercollegiate Faculty of Biotechnology University of Gdansk and Medical University of Gdansk, 2. ViroGenetics - BSL3 Laboratory of Virology, Malopolska Centre of Biotechnology, Jagiellonian University

44. Laboratory of Recombinant Vaccines, Intercollegiate Faculty of Biotechnology University of Gdansk and Medical University of Gdansk, 2. ViroGenetics - BSL3 Laboratory of Virology, Malopolska Centre of Biotechnology, Jagiellonian University

45. Laboratory of Recombinant Vaccines, Intercollegiate Faculty of Biotechnology University of Gdansk and Medical University of Gdansk, 2. ViroGenetics - BSL3 Laboratory of Virology, Malopolska Centre of Biotechnology, Jagiellonian University

46. Laboratory of Recombinant Vaccines, Intercollegiate Faculty of Biotechnology University of Gdansk and Medical University of Gdansk, 2. ViroGenetics - BSL3 Laboratory of Virology, Malopolska Centre of Biotechnology, Jagiellonian University

47. Laboratory of Recombinant Vaccines, Intercollegiate Faculty of Biotechnology University of Gdansk and Medical University of Gdansk, 2. ViroGenetics - BSL3 Laboratory of Virology, Malopolska Centre of Biotechnology, Jagiellonian University

48. Laboratory of Recombinant Vaccines, Intercollegiate Faculty of Biotechnology University of Gdansk and Medical University of Gdansk, 2. ViroGenetics - BSL3 Laboratory of Virology, Malopolska Centre of Biotechnology, Jagiellonian University

49. Laboratory of Recombinant Vaccines, Intercollegiate Faculty of Biotechnology University of Gdansk and Medical University of Gdansk, 2. ViroGenetics - BSL3 Laboratory of Virology, Malopolska Centre of Biotechnology, Jagiellonian University

5. Laboratory of Recombinant Vaccines, Intercollegiate Faculty of Biotechnology University of Gdansk and Medical University of Gdansk, 2. ViroGenetics - BSL3 Laboratory of Virology, Malopolska Centre of Biotechnology, Jagiellonian University

50. Laboratory of Recombinant Vaccines, Intercollegiate Faculty of Biotechnology University of Gdansk and Medical University of Gdansk, 2. ViroGenetics - BSL3 Laboratory of Virology, Malopolska Centre of Biotechnology, Jagiellonian University

51. Laboratory of Recombinant Vaccines, Intercollegiate Faculty of Biotechnology University of Gdansk and Medical University of Gdansk, 2. ViroGenetics - BSL3 Laboratory of Virology, Malopolska Centre of Biotechnology, Jagiellonian University

52. Laboratory of Recombinant Vaccines, Intercollegiate Faculty of Biotechnology University of Gdansk and Medical University of Gdansk, 2. ViroGenetics - BSL3 Laboratory of Virology, Malopolska Centre of Biotechnology, Jagiellonian University

53. Laboratory of Recombinant Vaccines, Intercollegiate Faculty of Biotechnology University of Gdansk and Medical University of Gdansk, 2. ViroGenetics - BSL3 Laboratory of Virology, Malopolska Centre of Biotechnology, Jagiellonian University

54. Laboratory of Recombinant Vaccines, Intercollegiate Faculty of Biotechnology University of Gdansk and Medical University of Gdansk, 2. ViroGenetics - BSL3 Laboratory of Virology, Malopolska Centre of Biotechnology, Jagiellonian University

56. Laboratory of Recombinant Vaccines, Intercollegiate Faculty of Biotechnology University of Gdansk and Medical University of Gdansk, 2. ViroGenetics - BSL3 Laboratory of Virology, Malopolska Centre of Biotechnology, Jagiellonian University

58. Laboratory of Recombinant Vaccines, Intercollegiate Faculty of Biotechnology University of Gdansk and Medical University of Gdansk, 2. ViroGenetics - BSL3 Laboratory of Virology, Malopolska Centre of Biotechnology, Jagiellonian University

59. Laboratory of Recombinant Vaccines, Intercollegiate Faculty of Biotechnology University of Gdansk and Medical University of Gdansk, 2. ViroGenetics - BSL3 Laboratory of Virology, Malopolska Centre of Biotechnology, Jagiellonian University

6. Laboratory of Recombinant Vaccines, Intercollegiate Faculty of Biotechnology University of Gdansk and Medical University of Gdansk, 2. ViroGenetics - BSL3 Laboratory of Virology, Malopolska Centre of Biotechnology, Jagiellonian University

60. Laboratory of Recombinant Vaccines, Intercollegiate Faculty of Biotechnology University of Gdansk and Medical University of Gdansk, 2. ViroGenetics - BSL3 Laboratory of Virology, Malopolska Centre of Biotechnology, Jagiellonian University

61. Laboratory of Recombinant Vaccines, Intercollegiate Faculty of Biotechnology University of Gdansk and Medical University of Gdansk, 2. ViroGenetics - BSL3 Laboratory of Virology, Malopolska Centre of Biotechnology, Jagiellonian University

62. Laboratory of Recombinant Vaccines, Intercollegiate Faculty of Biotechnology University of Gdansk and Medical University of Gdansk, 2. ViroGenetics - BSL3 Laboratory of Virology, Malopolska Centre of Biotechnology, Jagiellonian University

63. Laboratory of Recombinant Vaccines, Intercollegiate Faculty of Biotechnology University of Gdansk and Medical University of Gdansk, 2. ViroGenetics - BSL3 Laboratory of Virology, Malopolska Centre of Biotechnology, Jagiellonian University

7. Laboratory of Recombinant Vaccines, Intercollegiate Faculty of Biotechnology University of Gdansk and Medical University of Gdansk, 2. ViroGenetics - BSL3 Laboratory of Virology, Malopolska Centre of Biotechnology, Jagiellonian University

8. Laboratory of Recombinant Vaccines, Intercollegiate Faculty of Biotechnology University of Gdansk and Medical University of Gdansk, 2. ViroGenetics - BSL3 Laboratory of Virology, Malopolska Centre of Biotechnology, Jagiellonian University

9. Laboratory of Recombinant Vaccines, Intercollegiate Faculty of Biotechnology University of Gdansk and Medical University of Gdansk, 2. ViroGenetics - BSL3 Laboratory of Virology, Malopolska Centre of Biotechnology, Jagiellonian University

Aalborg University

Abbott

Abdaliyev Askar, Tungushbayev Talgat, Sharipova Saule, Shevtsov Alexandr, Amirgazin Asylulan, Kamalova Dinara, Ramankulov Erlan, Balykbaev Kanat

Academic Center for Pathomorphological and Genetic-Molecular Diagnostics ltd, Bialystok, Poland

Acibadem Mehmet Ali Aydinlar University School of Medicine, Medical Genetics Department

African Centre for Excellence for Genomics of Infectious Diseases (ACEGID), Redeemer’s University

African Centre of Excellence for Genomics of Infectious Diseases (ACEGID), Redeemer's University

African Centre of Excellence for Genomics of Infectious Diseases (ACEGID), Redeemer's University, Ede, Osun State, Nigeria

African Centre of Excellence for Genomics of Infectious Diseases (ACEGID), Redeemer’s University

African Centre of Excellence for Genomics of Infectious Diseases (ACEGID), Redeemer’s University, Ede

Agiomix

AIDS Vaccine Research Laboratories

Al-Quds Nutrition and Health Research Institute, Al-Quds University

Al Jalila Children’s Hospital

Al Jalila Genomics Center

Al Wathba Veterinary laboratory, Abu Dhabi Agriculture and Food Safety Authority (ADAFSA), Abu Dhabi, United Arab Emirates. PO Box 52150.

Alameda County Public Health Department

Alanagreh

Alaska State Virology Lab

Alaska State Virology Laboratory

Albert Einstein College of Medicine, Dept. of Microbiology & Immunology, Chandran lab

Alberta Precision Labs (APL)

Albertsen lab, Department of Chemistry and Bioscience, Aalborg University, Denmark

Albertsen Lab, Department of Chemistry and Bioscience, Aalborg University, Denmark

Alea Genetic Center

Alea Genetic Centre

Alea Geneticki Centar

Alsafar

Alsafar - Khalifa University Abu Dhabi

American Type Culture Collection Inc. (ATCC)

AMES Centro Polidiagnostico Strumentale S.r.l.

Andersen lab at Scripps Research

Andersen Lab, The Scripps Research Institute

ANOUAL

Area Biologia Molecolare - Istituto Zooprofilattico Sperimentale della Sicilia

Area Biologia Molecolare Istituto Zooprofilattico Sperimentale della Sicilia

Área de Secuenciación del Laboratorio de Virología del Hospital de Niños Dr. Ricardo Gutierrez on behalf of 'Proyecto Argentino Interinstitucional de genomica de SARS-CoV-2' (PAIS Consortium)

Area of Virology, Serology and Virology Division (SAViD), New South Wales Health Pathology Randwick

Area of Virology, Serology and Virology Division (SAVID), New South Wales Health Pathology Randwick

ARGO Laboratorio Genomica ed Epigenomica

ARGO Laboratorio Genomica ed EpigenomicA

ARGO Open Lab Platform for Genome sequencing

ARGO Open Lab Platform for Genome Sequencing

Arizona State Public Health Laboratory

Arizona State University

Army Medical and Veterinary Research Center

Army Medical Center, Scientific Department, Virology Laboratory

ARUP Laboratories

Associação Fundo de Incentivo a Pesquisa

Associação Fundo de Incentivo à Pesquisa (AFIP)

Associação Fundo de Incentivo à Pesquisa (AFIP).

Atlas Genomics

Av. Julián Coronel 905 entre Esmeraldas y José Mascote Av. Juan Tanca Marengo No. 100 y Av. de las Américas

AZ Delta

AZ Klina

AZ KLina

AZ SINT-JAN BRUGGE

AZDelta

Azerbaijan National Hematology Center Division of Medical Genetics

Bacteriology, Georgia Public Health Laboratory (GPHL)

Banc de Sang i Teixits

Bangladesh Council of Scientific and Industrial Research

Barts Health NHS Trust

Barts Health NHS TRust

Basic and Applied Research on Jute Project

Baylor College of Medicine/ GCID

Baylor College of Medicine: HGSC

Baylor Esoteric + Molecular Lab

Baylor Esoteric and Molecular Lab

Baylor Scott & White-Temple

Baylor Scott & White - Temple

BCCDC Public Health Laboratory

Beaconlab (Bioinformatics Evolution and Comparative Genomics lab), Dept of Biosciences, University of Milan

Beaconlab (Bioinformatics, Evolution and Comparative Genomics lab), Dept of Biosciences, University on Mila

Beaconlab (Bioinformatics, Evolution and Comparative Genomics lab), Dept of Biosciences, University on Milan

Beijing Center for Disease Prevention and Control

Beijing Genomics Institute (BGI)

Beijing Institute of Microbiology and Epidemiology

Bergthaler laboratory, CeMM Research Center for Molecular Medicine of the Austrian Academy of Sciences

Bezmialem Vakif University, Medical School & Beykoz Institute of Life Sciences & Biotechnology

BGI-shenzhen & The First Affiliated Hospital of Guangzhou Medical University

BGI & Institute of Microbiology, Chinese Academy of Sciences & Shandong First Medical University & Shandong Academy of Medical Sciences & General Hospital of Central Theater Command of People's Liberation Army of China

Bielefeld University

BIOBANCO / COCTI

Biobank Lab, Department of Molecular Biophysics, Faculty of Biology and Environmental Protection, University of Lodz

Biochemistry and Molecular Biology Department-Faculty of Medicine, Al-Quds University

Biocódices SA. on behalf of 'Proyecto Argentino Interinstitucional de genomica de SARS-CoV-2' (PAIS Consortium)

Biocruces

Biocruces-Bizkaia

Biocruces Bizkaia

BioCruces Bizkaia

BioInfo Experts, LLC

Bioinfoexperts, LLC

BioInfoExperts, LLC

Bioinformatics and Life Sciences, Noblis

Bioinformatics Division, National Institute of Biotechnology

Bioinformatics Division, National Institute of Biotechnology (NIB)

Bioinformatics Laboratory

Bioinformatics Laboratory - LNCC

Bioinformatics Laboratory / LNCC

Bioinformatics Research Group, Szentágothai Research Centre

Bioinformatics Research Group, Szentágothai Research Centre, University of Pécs

Bioinformatics, National institute of traditional medicine

Biolab Diagnostic Laboratories

Biologia molecular de enfermedades emergentes y EPOC, Instituto Nacional de Enfermedades Respiratorias

Biological prevention, army

Biological Prevention, Army

Biology, MCL

Biomedical Research Center (BRC)

Biomedical Sciences and Public Health, Polytechnic University of Marche

Bioptická laborator s.r.o.

Biosafety Department PCL3

Biosafety Level-3 Laboratory, Indonesian Institute of Sciences (LIPI)

Biotechnology & OMICs Laboratory

Biotechnology & OMICs Laboratory, Natural & Medical Sciences Research Center, University of Nizwa

Biotia

Botswana Harvard AIDS Institute Partnership

Botswana Harvard HIV Reference Laboratory

Botswana Harvard HIV Reference Laboratory,

Botswana Institute for Technology Research and innovation

Botswana Institute for Technology Research and Innovation

Breuer Lab, UCL

Brotman Baty Institute for Precision Medicine

BSL-3 Lab, National Institute for Viral Disease Control and Prevention, Chinese Center for Disease Control and Prevention

BSWMC-Temple Molecular

BTC, Khalifa University

Bundeswehr Institut of Microbiology

Bundeswehr Institute of Microbiology

Bureau of Public Health Laboratories, Florida Department of Health (BPHL, FLDOH)

Bursa Uludag University

Bushman Lab - University of Pennsylvania

Cadham Provincial laboratory

California Department of Public Health

Can Ruti SARS-CoV-2 Sequencing Hub (HUGTiP/IrsiCaixa/IGTP)

Cancer Biology Department, National Cancer Institute

Cantacuzino Institute

Cantacuzino Institute Virology

Carpi Laboratory - Purdue University

Carrington Lab, Department of PreClinical Sciences

Carrington Lab, Department of PreClinical Sciences, Building 36, First Floor Biochemistry Unit, Faculty of Medical Sciences, The University of the West Indies

Carrington Lab, Department of PreClinical Sciences, Faculty of Medical Sciences, The University of the West Indies

Carrington Lab, Department ofBuilding 36, First Floor Biochemistry Unit, Faculty of Medical Sciences, The University of the West Indies

CAS Key Laboratory of Special Pathogens and Biosafety and Center for Emerging Infectious Diseases

CBRN Defence and Security, Swedish Defence Research Agency

Cedars-Sinai Medical Center, Molecular Pathology Laboratory of Department of Pathology & Laboratory Medicine and Genomic Core

CEIRS Data Processing and Coordinating Center, Center for Research on Influenza Pathogenesis (CRIP)

CEIRS Data Processing and Coordinating Center, St. Jude Center of Excellence for Influenza Research and Surveillance (CEIRS)

CEITEC MU

Center for Biotechnology and Cell Therapy, São Rafael Hospital, Salvador, Brazil

Center for Genome Research and Biocomputing

Center for Genome Sciences, US Army Medical Research Institute of Infectious Disease (USAMRIID)

Center for Genomics and System Biology, New York University

Center for Global Health, University of New Mexico Health Sciences Center

Center for Laboratory Control of Infectious Diseases, Korea Centers for Diseases Control and Prevention

Center for Laboratory Medicine

Center for Laboratory Medicine St. Gallen

Center for Mathematical Modeling and Center for Genome Regulation. Santiago, Chile

Center for Medical Genetics, Keio University School of Medicine, Tokyo, Japan

Center for Personalized Medicine, Children's Hospital Los Angeles

Center for Research and Innovation, Faculty of Medical Technology, Mahidol University

Center for Virology

Center of Advanced Studies and Technology, CAST

Center of Advanced Studies and Technology, Molecular Genetics Laboratory

Center of Genomics and bioinformatics, Bioinformatics laboratory

Center of Medical Microbiology, Virology, and Hospital Hygiene, Heinrich Heine University Düsseldorf

Center of Medical Microbiology, Virology, and Hospital Hygiene, University of Duesseldorf

Center of Scientific Excellence for Influenza Viruses, National Research Centre (NRC), Egypt.

Center of Scientific Excellence for Influenza Viruses,National Research Centre (NRC), Egypt.

Centers for Disease Control and Prevention

Centers for Disease Control and Prevention Division of Viral Diseases, Pathogen Discovery

Centers for Disease Control and Prevention, Dengue Branch

Centers for Disease Control, R.O.C. (Taiwan)

Centogene

Centogene AG

Central Biological Research Laboratory and Department of Biochemistry and Molecular Biology

Central Biological Research Laboratory and Department of Biochemistry and Molecular Biology Central Biological Research Laboratory and Department of Biochemistry and Molecular Biology

Central Laboratories, Egyptian Ministry of Health and Population

Central Public Health Lab, National Public Health Organization

Central Public Health Laboratories, Egyptian Ministry of Health and Population

Central Public Health Laboratory - LACEN -Bahia, Salvador, Brazil

Central Virology Laboratory

Central Virology Laboratory, Israel Ministry of Health

Centre de Recherches Médicales de Lambaréné (CERMEL)

Centre For Biotechnology Research and Development

Centre for Clinical Infection and Diagnostics Research and Genomics Innovation Unit, Guy's and St. Thomas' NHS Trust

Centre for Dengue Research

Centre for Dengue Research and AICBU, Department of Immunology and Molecular Medicine

Centre for Dengue Research and AICBU, Department of Immunology and Molecular Medicine, University of Sri Jayewardenepura, Sri Lanka

Centre for Dengue Research, Department of Immunology and Molecular Medicine

Centre for Dengue Research, Department of Immunology and Molecular Medicine,

Centre for Dengue Research, USJ, SL

Centre for Human Virology & Genomics, Nigerian Institute of Medical Research

Centre Hospitalier Universitaire de Rouen Laboratoire de Virologie

Centre Muraz

Centre of Nanotechnologies, INCD IMT-Bucuresti (National Institute for Research and Development in Microtechnologies - Bucharest)

Centro Asistencial Docente y de Investigacion, Universidad de Magallanes

Centro de Desenvolvimento Tecnológico em Saúde - CDTS

Centro de Desenvolvimento Tecnologico em Saude, Fundacao Oswaldo Cruz

Centro de Investigación en Ciencias de la Salud y Biomedicina, U.A.S.L.P

Centro de Investigación en Ciencias de la Salud y Biomedicina, U.A.S.L.P.

Centro de Investigación en Ciencias de la Salud y Biomedicina, U.A.S.LP.

Centro de Investigaciones en Microbiología y Biotecnología-UR (CIMBIUR), Facultad de Ciencias Naturales, Universidad del Rosario, Bogotá, Colombia

Centro de Investigaciones en Microbiología y Biotecnología-UR (CIMBIUR), Facultad de Ciencias Naturales, Universidad del Rosario, Bogotá, Colombia Icahn School of Medicine at Mount Sinai, New York, USA

Centro de Investigaciones en Microbiología y Biotecnología-UR (CIMBIUR), Facultad de Ciencias Naturales, Universidad del Rosario, Bogotá, Colombia Instituto Nacional de Salud, Bogotá, Colombia Icahn School of Medicine at Mount Sinai, New York, USA

Centro de Secuenciación NASERTIC

Centro Polidiagnostico strumentale AMES

Cerba

Cerba Lab

Cerba LAb

CERBA LAB

Chan-Zuckerberg Biohub

Charité-Universitätsmedizin Berlin

Charite Universitaetsmedizin Berlin, Institute of Virology

Charité Universitätsmedizin Berlin, Institut für Virologie

Charité Universitätsmedizin Berlin, Institut für Virologie, Charitéplatz 1, 10117 Berlin, Germany

Charité Universitätsmedizin Berlin, Institut für Virologie/Labor Berlin

Charite Universitatsmedizin Berlin, Institute of Virology

Charité Universitätsmedizin Berlin, Institute of Virology

Charité Universitätsmedizin Berlin, Institute of Virology, Charitéplatz 1, 10117 Berlin, Germany

Charité Virology-University of Costa Rica

Child Health Research Foundation

Child Health Research Lab

Chinese PLA Institute for Disease Control and Prevention

Chiu Laboratory UCSF-Abbott Viral Diagnostics and Discovery Center University of California, San Francisco

Chiu Laboratory, University of California, San Francisco

Chongqing Municipal Center for Disease Control and Prevention

CHU Clermont-Ferrand, service de virologie

CHU LILLE

CHU Lille - Laboratoire de Virologie

CHU Nantes Virology

CHU NIMES

CHU Pontchaillou

CHU Purpan - Laboratoire de Virologie - Institut Fédératif de Biologie

CIAD LDM-LGM

Cicin-Sain Lab

Cicin Sain lab, Helmholtz Centre for Infection Research

CIDM-PH et al.

CIDM-PH, Westmead Hospital

CINVESTAV

CISLD (Clinical Institute of Special Laboratory Diagnostics), University Children's Hospital, University Medical Center Ljubljana

City of Milwaukee Health Department Laboratory

Clinical and Experimental Pharmacology Lab, LR16SP02, National Center of Pharmacovigilance, University of Tunis El Manar, Tunis, Tunisia. 2-Neurodegenerative diseases and psychiatric troubles, LR18SP03, Razi Hospital, University of Tunis El Manar, Tunis, Tunisia. 3- Ministry of Health, National Observatory of New and Emerging Diseases, 1006, Tunis, Tunisia

Clinical Center, University of Sarajevo; Unit for Clinical Microbiology

Clinical Diagnostics Laboratory, Diagnostic & Experimental Pathology, Lilly Research Laboratories

Clinical Division, Fred Hutchinson Cancer Research Center

Clinical Laboratory, Fuyang City Center for Disease Control and Prevention

Clinical Laboratory, Hospital Israelita Albert Einstein

Clinical Microbiology Lab RSPTN Universitas Hasanuddin

Clinical microbiology lab, RSPTN Universitas Hasanuddin

Clinical Microbiology Laboratory, Faculty of Medicine, Universitas Indonesia

Clinical microbiology, Sahlgrenska University Hospital

Clinical Microbiology, Sahlgrenska University Hospital,

Clinical Research Center, National Hospital Organization Nagoya Medical Center

Clinical virology

Clinical virology Laboratory, Stanford University School of Medicine

Clinical Virology Laboratory, Stanford University School or Medicine

CMBG FN Brno

CNR Virus des Infections Respiratoires - France SUD

Collaboration between the University of Melbourne at The Peter Doherty Institute for Infection and Immunity, and the Victorian Infectious Disease Reference Laboratory

Colleen B. Jonsson

College of Veterinary Medicine, Chungnam National University

Colorado Department of Public Health & Environment

Colorado Department of Public Health and Environment

Colorado Department of Puplic Health and Environment

Colorado State University - Ebel Lab

Communicable Disease Laboratory, Public Health Directorate

Compass Laboratory Services

Computational Virology Group, Center for Bacteria and Viruses Resources and Bioinformation, Wuhan Institute of Virology, Chinese Academy of Sciences,Wuhan 430071, China

Computer Science and Engineering

Contact:Hiroyuki Asakura Tokyo Metropolitan Institute of Public Health, Department of Microbiology

Contact:Ryota Kumagai Tokyo Metropolitan Institute of Public Health

Coordenação Geral de Laboratórios de Saúde Pública (CGLAB)

Coordenação Geral de Laboratórios de Saúde Pública (CGLAB/DAEVS/SVS/MS)

CoronaNet Lab- TaskForce Regione Campania, CEINGE Biotecnologie Avanzate, Via G. Salvatore

Corporacion Corpogen Universidad de los Andes Universidad Central

COVID-19 Genomics UK (COG-UK) Consortium

COVID-19 National Reference Laboratory

COVID-19 Network Investigations (CONI) Alliance

COVID Research Cell (CRC), Wazed Miah Science Research Center

Creighton University School of Medicine, Departments of Medical Microbiology and Pharmacology and Neuroscience

Croatian Institute of Public Health

Crosetto lab, Karolinska Institutet, SciLifeLab

Cruces University Hospital

CSIR-CDRI, Lucknow

CSIR-CDRI/SGPGI

CSIr-CDRI/SGPGI, Lucknow

CSIR-CDRi/SGPGI, Lucknow

CSIR-CDRI/SGPGI, Lucknow

CSIR-Centre for Cellular and Molecular Biology

CSIR-Centre for Cellular and Molecular Biology - INSACOG

CSIR-IGIB

CSIR-IGIB/Max

CSIR-Indian Institute of Chemical Biology, MEDICA Supercpecialty Hospital Kolkata

CSIR-Institute of Genomics and Integrative Biology

CSIR-Institute of Microbial Technology

CSIR-National Botanical Research Institute

CSIR Institute of Genomics and Integrative Biology

CSIR Institute of Genomics and Integrative Biology (CSIR-IGIB) / Max

CTMR, Karolinska Institutet, Stockholm, Sweden

CUB Hopital Erasme Laboratoire d'Anatomie Pathologique

Curative Inc

Cytocheck Laboratory

Dasman diabetes Institute

Dasman Diabetes Institute

Data Science

DC Public Health Lab/ Dept. of Forensic Sciences

deCODE genetics

Defence Research & Development Establishment

Defence Research & Development Establishment (DRDE)

Defence Services Medical Research Center, Biological Research Laboratory

Delaware Public Health Lab

Delaware Public Health Laboratory

Democritus University of Thrace, Department of Medicine

Dep. Of Oncology and Hemato-Oncology University of Milan

Department for Public Health Microbiology Ljubljana, National Laboratory for Health, Environment and Food

Department for Public Health Microbiology Ljubljana, NLZOH

Department for Virology, Molecular Biology and Genome Research, R. G. Lugar Center for Public Health Research, National Center for Disease Control and Public Health (NCDC) of Georgia.

Department of Acute Infectious Diseases Control and Prevention, Yunnan Provincial Center for Disease Control and Prevention

Department of Acute Infectious Diseases Control and Prevention,Yunnan Provincial Center for Disease Control and Prevention

Department of Biochemistry, Cell and Molecular Biology, West African Centre for Cell Biology of Infectious Pathogens (WACCBIP), University of Ghana

Department of Biology, University of Basrah

Department of Biomedical Informatics, University of Arkansas for Medical Sciences (UAMS)

Department of Biomedical, Surgical and Dental Sciences and Department of Biomedical Sciences for Health

Department of Biosystems Science and Engineering, ETH Zurich

Department of Biosystems Science and Engineering, ETH Zürich

Department of biotechnology, Yonsei University

Department of Clinical Diagnostics

Department of Clinical Laboratory, the First People's Hospital of Yunnan Province

Department of Developmental Medicine, Research Institute,

Department of Developmental Medicine, Research Institute, Osaka Women's and Children's Hospital

Department of Emerging Infectious Diseases, Institute of Tropical Medicine, Nagasaki University

Department of Epidemiology, Infectious Disease Control and Prevention, Hiroshima University, Japan

Department of Food safety, Nutrition and Veterinary public health - Istituto Superiore di Sanità

Department of General Diagnostics; Department of Virology; Istituto Zooprofilattico Sperimentale del Lazio e della Toscana (IZSLT)

Department of General Services Division of Consolidated Laboratory Services, Virginia Division of Consolidated Laboratory Services Sequencing Submission Group

Department of Health Technology and Informatics, Faculty of Health and Social Science, The Hong Kong Polytechnic University

Department of Health Technology and Informatics, Hong Kong Polytechnic University of Hong Kong

Department of Health Technology and Informatics, The Hong Kong Polytechnic University

Department of Healthcare Biotechnology, National University of Sciences and Technology (NUST)

Department of Immunology, The Scripps Research Institute

Department of Infection Prevention and Infectious Diseases, University Hospital Regensburg

Department of Infection, Immunity and Cardiovascular Disease, The Florey Institute, The Medical School, University of Sheffield

Department of Infectious Disease Prevention and Control, Henan Provincial Center for Disease Control and Prevention

Department of Infectious Diseases, Istituto Superiore di Sanità

Department of Infectious Diseases, Kobe Institute of Health

Department of Inspection,Centers for Disease Control and Prevention of Lishui

Department of Internal Medicine, College of Medicine, Chosun University

Department of Laboratory Medicine and Molecular Diagnostics, Sunnybrook Health Sciences Centre

Department of Laboratory Medicine Tan Tock Seng Hospital

Department of Laboratory Medicine, Lin-Kou Chang Gung Memorial Hospital, Taoyuan, Taiwan

Department of Laboratory Medicine, Lin-Kou Chang Gung Memorial Hospital, Taoyuan, Taiwan.

Department of Laboratory Medicine, Tan Tock Seng Hospital

Department of Laboratory, Medicine Tan Tock Seng Hospital

Department of Medical Laboratory Sciences, Arab American University

Department of Medical Microbiology

Department of Medical Microbiology, Faculty of Medicine, University of Malaya

Department of Medical Microbiology, Hospital Pengajar Universiti Putra Malaysia

Department of Medical Microbiology, Leiden University Medical Center

Department of Medical, Biotechnologies University of Siena

Department of Medicinal Genetics, Bursa Uludag University, Faculty of medicine By Sehime Gülsün Temel, Adem Alemdar, Kadir Yesilbag

Department of Medicine

Department of Microbial Biotechnology, Genetic Engineering Division, National Research Centre

Department of Microbial Biotechnology, Genetic Engineering Division, National Research Centre,

Department of Microbiology

Department of Microbiology and Immunology

Department of Microbiology and Immunology- SQUH

Department of Microbiology and Immunology-SQUH

Department of Microbiology and Immunology-SQUH Department of Microbiology and Immunology, Sultan Qaboos University Hopsital, P.O 35, Postal code 123

Department of Microbiology and Immunology - Pasteur Institute in Ho Chi Minh city

Department of microbiology laboratory,Anhui Provincial Center for Disease Control and Prevention

Department of Microbiology, College of Medicine and Medical Research Institute Chungbuk National University

Department of Microbiology, Faculty of Medicine, Chinese University of Hong Kong, Hong Kong SAR, China

Department of Microbiology, Gandhi Medical College and Hospital Secendrabad, Hyderbad, India

Department of Microbiology, Gandhi Medical College and Hospital, Secendrabad, Hyderabad

Department of Microbiology, Gandhi Medical College and Hospital, Secendrabad, Hyderabad, India

Department of Microbiology, Guangdong Provincial Center for Diseases Control and Prevention

Department of Microbiology, Institute for Viral Diseases, College of Medicine, Korea University

Department of Microbiology, PathWest QEII Medical Centre

Department of Microbiology, The University of Hong Kong

Department of Microbiology, University Hospital Motol

Department of Microbiology, Yokohama City University School of Medicine

Department of Microbiology, Zhejiang Provincial Center for Disease Control and Prevention

Department of Microbiology; Ryota Kumagai Tokyo Metropolitan Institute of Public Health

Department of Molecular & Medical Virology, Ruhr University Bochum, 44801 Bochum, Germany | Institute of Virology, Charité - Universitätsmedizin Berlin, corporate member of Freie Universität Berlin, Humboldt-Universität zu Berlin and Berlin Institute of Health (BIH), Berlin, Germany

Department of Molecular and Translational Medicine, Section of Microbiology, University of Brescia, ASST Spedali Civili, Brescia

Department of Molecular Medicine, Computational Medicine Group, Univeresity of Padova, Padova, Italy

Department of Molecular Medicine, University of Padova

Department of Molecular Medicine,Computational Medicine Group,Univeresity of Padova,Padova,Italy

Department of Molecular Virology, Cyprus Institute of Neurology and Genetics

Department of Neurovirology, National Institute of Mental Health and Neuroscience (NIMHANS)

Department of Neurovirology, National Institute of Mental Health and Neurosciences (NIMHANS)

Department of Omics Analysis

Department of Pathobiology, University of Connecticut

Department of Pathology and Laboratory Medicine, University of California Los Angeles

Department of Pathology and Medicine, New York University School of Medicine

Department of Pathology, National Institute of Infectious Diseases

Department of Public Health Microbiology Ljubljana, National Laboratory for Health, Environment and Food

Department of Respiratory & Other Viral Infections of L.V. Gromashevsky Institute of Epidemiology & Infectious Diseases NAMS of Ukraine, JSC "Farmak"

Department of Respiratory and Critical Care

Department of Respiratory and other Viral Infections of L.V.Gromashevsky Institute of Epidemiology & Infectious Diseases NAMS of Ukrainе, JSC "Farmak"

Department of Respiratory and other Viral Infections of L.V.Gromashevsky Institute of Epidemiology & Infectious Diseases NAMS of Ukraine, JSC "Farmak"

Department of Veterinary Biotechnalogy, College of Veterinary Science, Rajendranagar, PV Narsimha Rao Telengana Veterinary University

Department of Veterinary Pathology, University of Liege - FARAH

Department of Veterinary Science, National Institute of Infectious Diseases

Department of Virology

Department of Virology and Parasitology, Fujita Health University School of Medicin

Department of Virology and Parasitology, Fujita Health University School of Medicine

Department of Virology Faculty of Medicine, Medicum University of Helsinki

Department of Virology Institute of Tropical Medicine Nagasaki University

Department of Virology, Bangabandhu Sheikh Mujib Medical University

Department of Virology, Faculty of Medicine, University of Helsinki, Helsinki, Finland

Department of Virology, Faculty of Medicine, University of Helsinki, Helsinki, Finland generated and submitted to GISAID

Department of Virology, Henri Mondor University Hospital, Assistance Publique Hôpitaux de Paris, Université Paris-Est Créteil, INSERM U955

Department of Virology, Institute of Tropical Medicine, Nagasaki University, Nagasaki, Japan

Department of Virology, Pitié-Salpêtrière hospital

Department of Virology, Public Health Laboratories Division

Department of Virology, Public Health Laboratories Division, National Institute of Health

Department of Virology, University of Helsinki and Helsinki University Hospital, Helsinki, Finland

Departments of Pathology and Medicine, New York University School of Medicine

Dept. Food safety, Nutrition and Veterinary Public Health, Istituto superiore di sanità

Dept. of Laboratory Medicine

Dept. of Microbiology and Infection Control, Akershus University Hospital HF

Dept. OPA, Beijing Institute of Microbiology and Epidemiology

DeRisi Lab, University of California, San Francisco

Diagen

Diagnostic- and Research Institute of Pathology, Medical University of Graz

Diagnostic and Research Center of Infectious Diseases, Medical Faculty, Andalas University

Diagnostic Genomics Lab and Functional Genomics Core, University of South Carolina

Diagnostic Genomics Laboratory

Diagnostic Virology Laboratory, National Veterinary Services Laboratories, USDA1920 Dayton Avenue, Ames, IA 50010, USA

Diagnostic Virology Laboratory, United States Department of Agriculture, National Veterinary Services Laboratories

Diagnostic Virology Laboratory, United States Department of Agriculture, National Veterinary Services Laboratories, Population Medicine and Diagnostic Sciences, Cornell University

Diagnostic Virology Laboratory, USDA National Veterinary Services Laboratories

Diagtron Laboratoria

Diarrhea department, National Institute for Viral Disease Control and Prevention, China CDC

Díaz-Muñoz Lab, Department of Microbiology and Molecular Genetics, University of California, Davis

Dipartimento di Biotecnologie Mediche

Dipartimento di Biotecnologie Mediche, University of Siena

Dipartimento di Scenze Biomediche e Cliniche, L. Sacco, Università di Milano

Dipartimento di Scienze Biomediche e Cliniche, L.Sacco, Università di Milano

Dirk Dittmer

Discovery DNA

Division of Emerging Infectious Diseases, Bureau of Infectious Diseases Diagnosis Control, Korea Disease Control and Prevention Agency

Division of Medical Virology, Stellenbosch University and National Health Laboratory Service (NHLS)

Division of Medical Virology, Stellenbosch University and NHLS Tygerberg Hospital

Division of Pathogen Resource Management, Korea National Institute of Health, Korea Disease Control and Prevention Agency

Division of Viral Diseases, CDC Pathogen Discovery Team

Division of Viral Diseases, Center for Laboratory Control of Infectious Diseases, Korea Centers for Diseases Control and Prevention

Division of Viral Diseases, Centers for Disease Control and Prevention

DMR

DMR_Myanmar

DNA Sequencing and Synthesis Facility (oligo.pl), Institute of Biochemistry and Biophysics PAS

DNA Solution Ltd

DNA Solution Ltd.

DNA SOLUTION LTD.

DNA Solution Ltd. L-5

DPH, Massachusetts State Public Health Lab

Dr. Jeff Wrana, Senior Investigator

Dr. Qudrat-I-Khuda Road, Dhaka-1205, Bangladesh

Dr. Thomas Vanderford's Lab

DRESDEN-concept Genome Center, CMCB, TU Dresden

Egyptian National Cancer Institute (ENCI)

Eijkman Institute for Molecular Biology, Ministry of Research and Technology/National Agency for Research and Innovation

Eijkman Institute for Molecular Biology, Ministry of Research and Technology/National Agency for Research and Innovation; National Institute of Health Research and Development

Enteric Viruses Group, ICMR-National Institute of Virology

Environmental and Global Health

Environmental and Global Health, University of Florida

Epiclin

Epigenetics, Saarland University

Erasmus Medical Center

Etlik Veterinary Control Central Research Institute

Evandro Chagas Institute

Evandro Chagas Institute Virology

Expert Microbiology, National Institute for Health and Welfare

Facultad de Ciencias (Sección Genética Evolutiva, Sección Virología).

Facultad de Ciencias de la Vida, UNAB

Faculty of Medicine

Faculty of Medicine, Al-Quds University

Faculty of Medicine, Universitas Indonesia

Faculty of Medicine, Universitas Sumatera Utara; Institute of Tropical Disease, Universitas Airlangga

Faculty of Natural Sciences, Comenius University in Bratislava

Faculty of Natural Sciences, Comenius University, Bratislava

Faroese National Reference Laboratory for Fish and Animal Diseases

Federal Budget Institution of Science, State Research Center for Applied Microbiology & Biotechnology

Fimlab Laboratories

Fimlab Laboratories, Arvo Ylpön katu 4, 33520 Tampere, Finland

Florida Bureau of Public Health Laboratories

Florida Bureau of Public Health Laboratories, Florida Department of Health

França Lab

Fred Hutchinson Cancer Research Center

Friedrich-Loeffler-Institut, Laboratory for NGS and Microarray Diagnostics

FSBSI "Chumakov Federal Scientific Center for Research and Development of Immune-and-Biological Products of Russian Academy of Sciences" & NRC "Kurchatov institute"

Fujian Center for Disease Control and Prevention

Fujita Health University School of Medicine, Department of Microbiology

Fujita health University, Department of Microbiology

Fujita health University, School of Medicine, Department of Microbiology

Fukuoka Institute of Health and Environmental Sciences

Fulgent Genetics

Functional Genomic Platform UATRS-biology, CNRST

Functional Genomic Platform/Service Analyses Biologique/UATRS/ Centre National Pour la Recherche Scientifique Et Technique (CNRST)

Functional Genomics Core, Center For Targeted Therapeutics,

Functional Genomics Core, University of South Carolina

Functional Genomics Core, University of South Carolina,

G42 Healthcare

G5 Evolutionary Genomics of RNA viruses, Virology Department, Institut Pasteur

GA Department of Public Health

Gagnon Lab, Southern Illinois University

Gazi University Faculty of Medicine, Medical Virology Laboratory

Geelong Centre for Emerging Infectious Diseases

Gen Era Diagnostics Corporation Life Science Research and Molecular Diagnostics

Gen Era Diagnostics Inc.

GenBio

Gencore- Universidad de los Andes

Gencore - Universidad de los Andes

Genelabs Medical (Pvt) Ltd

Genetica y Virologia, Facultad de Ciencias

Genetics Research Center, University of Social Welfare and Rehabilitation Sciences

Genetics Research Center. University Of Social Welfare And Rehabilitation Sciences

Genetics Working Group (Pokja Genetik) Faculty of Medicine, Public Health and Nursing Universitas Gadjah Mada (FK-KMK UGM), Disease Investigation Center Wates Ministry of Agriculture Indonesia, Department of Microbiology FK-KMK UGM, Laboratorium Diagnostik Yayasan Tahija World Mosquito Program (WMP) Yogyakarta Center for Tropical Medicine FK-KMK UGM, Integrated Research Center FK-KMK UGM, Department of Computer Science and Electronics FMIPA UGM

Genetics Working Group (Pokja Genetik) Faculty of Medicine, Public Health and Nursing Universitas Gadjah Mada (FK-KMK UGM); Disease Investigation Center Wates Ministry of Agriculture Indonesia; Department of Microbiology FK-KMK UGM; Laboratorium Diagnostik Yayasan Tahija World Mosquito Program (WMP) Yogyakarta Center for Tropical Medicine FK-KMK UGM; Integrated Research center FK-KMK UGM; Department of Computer Science and Electronics FMIPA UGM

Genetics Working Group (Pokja Genetik) Faculty of Medicine, Public Health and Nursing Universitas Gadjah Mada (FK-KMK UGM); Disease Investigation Center Wates Ministry of Agriculture Indonesia; Department of Microbiology FK-KMK UGM; Laboratorium Diagnostik Yayasan Tahija World Mosquito Program (WMP) Yogyakarta Center for Tropical Medicine FK-KMK UGM; Integrated Research Center FK-KMK UGM; Department of Computer Science and Electronics FMIPA UGM

Genetics Working Group (Pokja Genetik) Faculty of Medicine, Public Health and Nursing Universitas Gadjah Mada (FK-KMK UGM); Disease Investigation Center Wates Ministry of Agriculture Indonesia; Department of Microbiology FK-KMK UGM; Laboratorium Diagnostik Yayasan Tahija World Mosquito Program (WMP) Yogyakarta Center for Tropical Medicine FK-KMK UGM; Integrated Research Center FK-KMK UGM; Department of Computer Science and Electronics FMIPA UGM; Balai Besar Teknik Kesehatan Lingkungan dan Pengendalian Penyakit (BBTKLPP) Yogyakarta

Genetics Working Group (Pokja Genetik) Faculty of Medicine, Public Health and Nursing Universitas Gadjah Mada (FK-KMK UGM); Disease Investigation Center Wates Ministry of Agriculture Indonesia; Department of Microbiology FK-KMK UGM; Laboratorium Diagnostik Yayasan Tahija World Mosquito Program (WMP) Yogyakarta Center for Tropical Medicine FK-KMK UGM; Integrated Research Center FK-KMK UGM; Department of Computer Science and Electronics FMIPA UGM; RSUP Dr. Sardjito

GENOMA MAYOR

Genome Analysis Center, Kamma Memorial Hospital

Genome Analysis Center, Yamanashi Central Hospital

Genome Center

Genomes & Disease, Center for Research in Molecular Medicine and Chronic Diseases, University of Santiago de Compostela

Genomic Laboratory (GLAB), Istanbul Technical University

Genomic platform

Genomic Research Lab, BCSIR

Genomic Research Lab, BCSIR, Dr. Qudrat-E-Khuda Road, Dhaka-1205, Bangladesh

Genomic Research Lab, BCSIR, Dr. Qudrat-E-Khuda Road, Dhaka 1205, Bangladesh

Genomic Research Lab. BCSIR

Genomic Sciences, Rehman Medical Institute

Genomics and Computational Biology Lab, Scientific Research Institute of Physical-Chemical Medicine, FMBA of Russia

Genomics and Discovery, Respiratory Viruses Branch, Division of Viral Diseases, Centers for Disease Control and Prevention

Genomics and Transcriptomics, Philip Morris International

Genomics Lab NIBD

Genomics Program, Children Cancer Hospital

genXone SA, Research & Development Laboratory

genXone SA, Research & Development Laboratory; The Faculty of Mathematics, Informatics and Mechanics of the University of Warsaw

GHC Genetics, s.r.o.

GIGA Medical Genomics

Ginkgo Bioworks Clinical Laboratory

Goethe University Hospital Frankfurt, Institute for Medical Virology

Gonoshasthaya-RNA Research Center, Gonoshasthaya-RNA Molecular Diagnostics and Research Center

Gonoshasthya-RNA Molecular Diagnostic and Research Center

Gonoshasthya-RNA Molecular Research Center

Gorgas memorial Institute for Health Studies

Gorgas memorial Institute For Health Studies

Gorgas Memorial Institute for Health Studies

Gorgas Memorial Laboratory of Health Studies

GRAM2.0, Université de Caen Normandie

GRAM2.0, Université de Caen Normandie / Laboratoire de Virologie, CHU de Caen / National Reference Center for Respiratory Viruses France Nord, Institut Pasteur, Paris. / Laboratoire de la rage et de la faune sauvage ANSES, Malzéville.

GRAM2.0, Université de Caen Normandie Laboratoire de Virologie, CHU de Caen

Greek Genome Center, Biomedical Research Foundation of the Academy of Athens (BRFAA)

Group for molecular genetics of pathogens

Group of Genomics and Postgenomic Technologies of Central Research Institute of Epidemilology

Group of Genomics and Postgenomic Technologies of Central Research Institute of Epidemiology

Grubaugh Lab - Yale School of Public Health

Grupo de Genómica y Bioinformática del Instituto de Investigación de la Cadena Láctea CONICET-INTA on behalf of 'Proyecto Argentino Interinstitucional de genomica de SARS-CoV-2' (PAIS Consortium)

Grupo de Investigaciones Microbiológicas-UR (GIMUR), Departamento de Biología, Facultad de Ciencias Naturales, Universidad del Rosario, Bogotá, Colombia Instituto Nacional de Salud, Bogotá, Colombia Icahn School of Medicine at Mount Sinai, New York, USA

Guangdong Provincial Center for Disease Control and Prevention

Guangdong Provincial Center for Diseases Control and Prevention

Guangdong Provincial Institution of Public Health

Gujarat Biotechnology Research Centre

Gujarat Biotechnology Research Centre, Gandhinagar

Gunma Prefectural Institute of Public Health and Environmental Sciences

Hangzhou Center for Disease and Control Microbiology Lab

Hangzhou Center for Disease Control and Prevention

Hangzhou Center for Diseases Control and Prevention

Hannover Medical School, Institute of Virology

HCRC

Health & Environment Institute of Gwangju

Health and Environmental Research Institute of Gwangju Metropolitan city

Health and Environmental Testing Laboratory

Hebei Provincial Center for Disease Control and Prevention, Shijiazhuang, Hebei Province; National Institute for Viral Disease Control and Prevention, China CDC

HEGP - Laboratoire de Virologie

Heinrich Pette Institute, Leibniz Institute for Experimental Virology

Helix

Hellenic Pasteur Institute, National Influenza Reference laboratory of Southern Greece & Unit of Bioinformatics and Applied Genomics

Hellenic Pasteur Institute, Public Health Laboratories, Unit of Bioinformatics and Applied Genomics

Hematopathology Laboratory, ACTREC, TMC

Héritas

HIV Molecular Lab

HIV molecular lab, Ethiopian Public Health Institute, Ethiopian

HIV Molecular Laboratory

HIV Molecular Laboratory, Ethiopian Public Health Institute

Hokkaido Institute of Public Health

Hong Kong Children's Hospital

Hong Kong Children’s Hospital

Hong Kong Department of Health

Hôpital Henri-Mondor Ap-Hp

Hospital General Universitario Gregorio Marañón

Hospital Ramón y Cajal

Hospital Regional Ushuaia - Centro Austral De Investigaciones Científicas - Universidad Nacional De Tierra Del Fuego on behalf of 'Proyecto Argentino Interinstitucional de genomica de SARS-CoV-2' (PAIS Consortium)

Hospital São Rafael - IDOR

Hospital Universitari Vall d'Hebron - Vall d'Hebron Institut de Recerca

Hospital Universitari Vall d'Hebron - Vall Hebron Institut de Recerca

Hospital Universitario 12 de Octubre

Hospital Universitario La Paz

Houston Health Department

Houston Health Department, Disease Prevention and Control

Houston Health Dept.

Houston Methodist Hospital

Hubei Provincial Center for Disease Control and Prevention

Hudsonalpha Genome Sequencing Center

HUG, Laboratory of Virology and the Health2030 Genome Center

HUG, Laboratory of Virology and Universitätsspital Basel

Human Genetic Research Center, Kawsar Biotech Company

Human Genome Center

Human Genome Variation Research Group, Malopolska Centre of Biotechnology

Hyde Lab

IAME UMR1137 Inserm, Université de Paris, Hôpital Bichat

Ibaraki Prefectural Institute of Public Health

IBL

IBP-laboratoire de virologie

ICAR-National Institute of High Security Animal Diseases

ICMR-NATIONAL INSTITUTE OF VIROLOGY, MICROBIAL CONTAINMENT COMPLEX

ID Genomics

Idaho Bureau of Laboratories

IEH and ID Genomics

IHU Mediterranee Infection

IHU Méditerranée Infection

IIP Institute of Genomics and Integrative Biology

IKOM, NTNU

ILBS - IGIB

Illinois Department of Public Health - Chicago Lab

Illinois Department of Public Health - Springfield Lab

Illumina Miseq

Imelda Hospital

Imelda hospital Bonheiden

Immunogenomics group, Institute of Life Sciences, Bhubaneswar

Immunogenomics lab, Institute of Life Sciences, Bhubaneswar

Immunology, Noguchi Memorial Institute for Medical Research

Imperial College London

IN State Department of Health Laboratory Services

Inciensa, Instituto Costarricense de Investigación y Enseñanza en Nutrición y Salud

Incubadora Venezolana de Ciencia, Venezuela / Instituto Nacional de Salud, Bogotá, Colombia / Grupo de Investigaciones Microbiológicas-UR (GIMUR), Departamento de Biología, Facultad de Ciencias Naturales, Universidad del Rosario, Bogotá, Colombia / Icahn School of Medicine at Mount Sinai, New York, USA

Indian Council of Medical Research-National Institute of Virology, Maximum Containment Laboratory

Indian Council of Medical Research-National Institute of Virology, Microbial Containment Complex

Indian Council of Medical Research-National Institute of Virology,Maximum Containment Laboratory

Infection and Immunology, Translational Health Science and Technology Institute

Infectious Disease Biology, Institute of Life Sciences

Infectious Disease Control and Prevention Institute

Infectious Disease Control Center, Center for Disease Control and Prevention of PLA

Infectious Disease Program, Broad Institute of Harvard and MIT

Infectious Disease Research Department, King Abdullah International Medical Research Center (KAIMRC)

Infectious Diseases Research, King Abdullah International Medical Research Center (KAIMRC)

Infectious Diseases, Chan-Zuckerberg Biohub Covidtracker

Infectious diseases, Chan Zuckerberg Biohub

Infectious Diseases, NC SLPH COVID-19 Response Team

Infectious Diseases, North Carolina State Laboratory of Public Health COVID-19 Response Team

Infectious Diseases, Quest Diagnostics

Influenza Virus Research Center, National Institute of Infectious Diseases

INMI Lazzaro Spallanzani IRCCS

Innovative Genomics Institute, UC Berkeley

Innovative Genomics Institute, UCB

INSACOG-KA, NIMHANS

Insepction Center of Hangzhou Center for Disease Control and Prevention

Inspection Center of Hanghzou Center for Disease Control and Prevention

Inspection Center of Hangzhou Center for Disease Control and Prevention

INSPI-Centro de Investigación Multidisciplinaria de la DTIDI

INSPI - Charité

Institut de Pathologie et Genetique

Institut de Pathologie et Génétique

Institut de Pathologie et Genetique (IPG)

Institut de Pathologie et Génétique (IPG)

Institut für Medizinische Virologie, Universitätsklinikum Frankfurt

Institut of Human Genetics, University Medicine Goettingen

Institut Pasteur CIBU-ERI

Institut Pasteur CIBU / ERI

Institut Pasteur CIBU /ERI

Institut Pasteur de Dakar

Institut Pasteur de la Guadeloupe

Institut Pasteur de la Guyane

Institut Pasteur de Montevideo

Institut Pasteur du Maroc

Institut Pasteur, Laboratory for Urgent Response to biological Threats

Institute for Computational Biomedicine, Weill Cornell Medicine

Institute for Developing Science and Health Initiatives

Institute for Developing Science and Health Initiatives (ideSHi)

Institute for Infectious Diseases, University of Bern

Institute for Infectious Diseases, University of Bern, Switzerland

Institute for infectious medicine & hospital hygiene, CaSe-Group

Institute for Medical Research Infectious Disease Research Centre, National Institutes of Health, Ministry of Health Malaysia

Institute for Medical Research, Infectious Disease Research Centre, National Institutes of Health, Minis

Institute for Medical Research, Infectious Disease Research Centre, National Institutes of Health, Ministry of Health Malaysia

Institute for Medical Virology, Goethe University Hospital Frankfurt

Institute for Systems Biology

Institute for Virology, University Hospital Duesseldorf, Medical Faculty, Heinrich-Heine-University Duesseldorf

Institute information KU Leuven, Clinical and Epidemiological Virology

Institute of Applied Biotechnologies a.s.

Institute of Applied Genomics

Institute of Biomedicine (IBiMED), Universidade de Aveiro

Institute of Biotechnology, DNA Sequencing and Genomics Laboratory, University of Helsinki

Institute of Biotechnology, Life Sciences Center, Vilnius University

Institute of Biotechnology, Life Sciences Center, Vilnius University and Thermo Fisher Scientific

Institute of Clinical Microbiology and Hygiene, University Hospital Regensburg

Institute of Disease Control and Prevention, People's Liberation Army

Institute of Environmental Science and Research (ESR)

Institute of Genomics and Integrative Biology - Council of Scientific and Industrial Research

Institute of Genomics Core Facility, University of Tartu

Institute of Health and Community Medicine

Institute of Human Genetics, Polish Academy of Sciences

Institute of Human Genetics, Polish Academy of Sciences,

Institute of Human Genetics, University Medical Center Goettingen

Institute of Human Virology, Zhongshan School of Medicine, Sun Yat-sen University

Institute of infectious medicine & hospital hygiene, CaSe-Group

Institute of Life Sciences

Institute of Life Sciences - INSACOG

Institute of Medical Biology, Chinese Academy of Medical Sciences and Peking Union Medical College

Institute of Medical Genetics and Applied Genomics

Institute of medical Microbiology and hospital Hygiene

Institute of Medical Microbiology and Hospital Hygiene

Institute of Medical Virology

Institute of Medical Virology, University of Zurich

Institute of microbiology and Immunology, Faculty of Medicine, University of Belgrade

Institute of Microbiology and Immunology, Faculty of Medicine, University of Ljubljana

Institute of Microbiology Universidad San Francisco de Quito

Institute of Microbiology, Chinese Academy of Sciences

Institute of Microbiology, Universidad San Francisco de Quito

Institute of Microbiology, University of Veterinary and Animal sciences

Institute of Molecular and Translational Medicine / Laboratory of Experimental Medicine

Institute of Molecular and Translational Medicine / Laboratory of Experimental Medicine, Faculty of Medicine and Dentistry, Palacky University

Institute of Molecular Biology and Biotechnology, National Institutes of Health

Institute of Molecular Biology NAS RA, Republic of Armenia, Department of Bioengineering, BioinformaticsInstitute and Molecular Biology IBMPh RAU, Republic of Armenia

Institute of Molecular Medicine, Section for Molecular Cell Biology

Institute of Molecular Virology, University Münster

Institute of Pathogen Biology, Chinese Academy of Medical Sciences & Peking Union Medical College

Institute of pathogenic microbiology, Jiangsu Provincial Center for Disease Control and Prevention

Institute of Public Health of Republic of North Macedonia Laboratory of Virology and Molecular Diagnostics

Institute of Tropical Disease, Universitas Airlangga

Institute of Tropical Disease, Universitas Airlangga; Faculty of Medicine, Universitas Sumatera Utara

Institute of Tropical Disease, Universitas Airlangga; Fakultas Kedokteran, Universitas Sumatra Utara

Institute of Tropical Medicine

Institute of Viral Disease Control and Prevention, China CDC

Institute of Virology, Biomedical Research Center of the Slovak Academy of Sciences, Bratislava; Comenius University Science Park, Bratislava

Institute of Virology, Charité – Universitätsmedizin Berlin

Institute of Virology, Clinial Virus Genomics, Medical Center, University of Freiburg, Freiburg, Germany

Institute of Virology, University Hospital, University of Bonn and German Center for Infection Research (DZIF), Bonn-Cologne, Bonn, Germany

Institute of Virology, University of Cologne

Instituto Adolfo Lutz Interdisciplinary Procedures Center Strategic Laboratory

Instituto Adolfo Lutz, Interdiciplinary Procedures Center, Strategic Laboratory

Instituto Adolfo Lutz, Interdisciplinary Procedures Center, Strategic Laboratory

Instituto Butantan (genome assembly and bioinformatics) and Mendelics (sequencing)

Instituto Butantan / Mendelics

Instituto de Biotecnologia - UNESP-Botucatu-SP

Instituto de Biotecnología de la UNAM

Instituto de Biotecnología, IABIMO (CONICET), Instituto de Virología, IVIT(CONICET), Instituto de Patobiología, IPVET(CONICET), CICVyA, INTA on behalf of 'Proyecto Argentino Interinstitucional de genomica de SARS-CoV-2' (PAIS Consortium)

Instituto de Diagnostico y Referencia Epidemiologicos

Instituto de Diagnóstico y Referencia Epidemiologicos

Instituto de Diagnóstico y Referencia Epidemiológicos

Instituto de Diagnostico y Referencia Epidemiologicos (INDRE)

Instituto de diagnóstico y Referencia Epidemiologicos (INDRE)

Instituto de diagnóstico y Referencia Epidemiologicos (INDRE) Departamento de Virología

Instituto de Medicina Tropical da Univesidade de São Paulo

Instituto de Medicina Tropical de Sao Paulo

Instituto de Medicina Tropical Universidade de São Paulo

Instituto de Patologia Vegetal (CIAP-INTA) on behalf of 'Proyecto Argentino Interinstitucional de genomica de SARS-CoV-2' (PAIS Consortium)

Instituto de Salud Carlos III

Instituto de Salud Publica de Chile

Instituto Gulbenkian de Ciencia

Instituto Gulbenkian de Ciência

Instituto Nacional de Ciencias Medicas y Nutricion

Instituto Nacional de Ciencias Medicas y Nutricion Infectious Diseases

Instituto Nacional de Ciencias Medicas y Nutricion Salvador Zubiran

Instituto Nacional de Enfermedades Respiratorias

Instituto Nacional de Enfermedades Respiratorias (INER)

Instituto Nacional de Enfermedades Respiratorias (INER), Centro de Investigación en Enfermedades Infecciosas (CIENI)

Instituto Nacional de Enfermedades Respiratorias (INER): Centro de Investigación en Enfermedades Infecciosas (CIENI)

Instituto Nacional de Investigación en Salud Publica

Instituto Nacional de Investigación en Salud Pública

Instituto Nacional de Investigación y Tecnología Agraria y Alimentaria (INIA)/ Departamento de Mejora Genética Animal

Instituto Nacional de Medicina Genomica

Instituto Nacional de Medicina Genómica

Instituto Nacional de Salud

Instituto Nacional de Salud- Direccion de Investigacion en Salud Publica

Instituto Nacional de Salud- Dirección de Investigación en Salud Pública

Instituto Nacional de Salud- Dirección de Investigación en Salud Pública, Universidad de los Andes- Applied genomics research group, Vicerrectoria de Investigación y Creación, Universidad de los Andes- Systems and Computing Engineering Department

Instituto Nacional de Salud- Dirección de Investigación en Salud Pública, Universidad de los Andes- Gencore

Instituto Nacional de Salud - Dirección de Investigación en Salud Pública

Instituto Nacional de Salud - Unidad de Secuenciación y Genómica

Instituto Nacional de Salud Universidad Cooperativa de Colombia Instituto Alexander von Humboldt Imperial College-London London School of Hygiene & Tropical Medicine

Instituto Nacional de Salud, Bogotá, Colombia

Instituto Nacional de Salud, Universidad Cooperativa de Colombia, Instituto Alexander von Humboldt, Imperial College-London, London School of Hygiene & Tropical Medicine

Instituto Nacional de Saude (INSA)

Instituto Nacional de Saude (INSA) and BioSystems & Integrative Sciences Institute (BioISI) Genomics Unit, FCUL

Instituto Nacional de Saude (INSA) and i3S - Instituto de Investigação e Inovação em Saúde

Instituto Nacional de Saude (INSA) and Institute of Biomedicine (iBiMed), Universidade de Aveiro

Instituto Nacional de Saude (INSA) and Instituto Gulbenkian de Ciencia (IGC)

Instituto Nacional Enfermedades Infecciosas C.G.Malbran

Instituto Octávio Magalhães / Fundação Ezequiel Dias (IOM/Funed)

INT Fondazione Pascale

Integrative Pharmacogenomics Institute (iPROMISE)

International Centre for Diarrhoeal Disease Research (ICDDR,B)

International Centre for Genetic Engineering and Biotechnology (ICGEB) and ARGO Open Lab Platform

International Centre for Genetic Engineering and Biotechnology (ICGEB) and ARGO Open Lab Platform for Genome Sequencing

International Centre for Genetic Engineering and Biotechnology (ICGEB)and ARGO Open Lab Platform

IRBA, 2MI

IRCCS Regina Elena National Cancer Institute

IRESSEF

IRESSEF GENOMICS LAB

Irish Coronavirus Sequencing Consortium-Teagasc Grange

Irish Coronavirus Sequencing Consortium - Helixworks

Irish Coronavirus Sequencing Consortium - National University of Ireland Galway

Irish Coronavirus Sequencing Consortium - National Virus Reference Laboratory

Irish Coronavirus Sequencing Consortium - Teagasc Moorepark

Irish Coronavirus Sequencing Consortium - Teagasc Oakpark

IrsiCaixa

IrsiCaixa - Can Ruti CovidSeq

IrsiCaixa AIDS Research Lab

IrsiCaixa Retrovirology Lab

Israel Central Virology laboratory

Israel Institute for Biological Research

Israel National Consortium for SARS-CoV-2 sequencing

Istanbul University-Cerrahpasa, Cerrahpasa School of Medicine, COVID-19 Laboratory

Istitituto Zooprofilattico Sperimentale dell'Abruzzo e Molise "G.Caporale"

Istituto di Genomica Applicata

Istituto di Genomica Applicata; Istituto Zooprofilattico Sperimentale delle Venezie

Istituto Nazionale Malattie Infettive Lazzaro Spallanzani IRCCS

Istituto Superiore di Sanità

Istituto Superiore di Sanità (ISS)

Istituto Zooprofilattico Sperimentale del Mezzogiorno

Istituto Zooprofilattico Sperimentale dell'Abruzzo e del Molise "G. Caporale".

Istituto Zooprofilattico Sperimentale dell'Abruzzo e Molise "G. Caporale"

Istituto Zooprofilattico Sperimentale dell'Abruzzo e Molise "G.Caporale"

Istituto Zooprofilattico Sperimentale dell’Abruzzo e Molise “G. Caporale”

Istituto Zooprofilattico Sperimentale della Lombardia e dell'Emilia Romagna (IZSLER), Risk Analysis and Genomic Epidemiology Unit

Istituto Zooprofilattico Sperimentale della Puglia e della Basilicata

Istituto Zooprofilattico Sperimentale della Sicilia

Istituto Zooprofilattico Sperimentale delle Venezie

IU-Cerrahpasa, Cerrahpasa School of Medicine, COVID-19 Lab

IZSM

James Molecular Lab

James Molecular Lab - OSUWMC

James Molecular Laboratory

James Polaris Molecular Laboratory

Jamil-ur-Rahman Center for Genome Research, Dr. Panjwani Center for Molecular Medicine and Drug Research

Jamil-ur-Rahman Center for Genome Research, Dr. Panjwani Center for Molecular Medicine and Drug Research, International Center for Chemical and Biological Sciences, University of Karachi

Jeremy Wang

Jessa

Jiangsu Provincial Center for Disease Control & Prevention

Jiangxi province Center for Disease Control and Prevention

Jiangxi Province Center for Disease Control and Prevention

Johns Hopkins Hospital Department of Pathology

Jonathan Li laboratory

Jonathan Li Laboratory

Kabara Cancer Research Institute

Kafkas University, Faculty of Medicine, Department of Medical Microbiology

Kansas Health and Environmental Lab

Kansas State University Veterinary Diagnostic Laboratory

Kansas State Veterinary Diagnostic Laboratory

Karolinska University Hospital

Kashi Clinical Laboratory

Kawsar Human Genetic Research Center

Kawsar Human Genetic Research Company

Keio University Hospital

Keio University School of Medicine

KEMRI-Wellcome Trust Research Programme,Kilifi

KEMRI-Wellcome Trust Research Programme/KEMRI-CGMR-C Kilifi

Kenema Government Hospital, Ministry of Health and Sanitation

Kentucky State Public Health Lab

Key Laboratory of Human Diseases, Comparative Medicine, Institute of Laboratory Animal Science

Key Laboratory of Medical Molecular Virology (MOE/NHC/CAMS), School of Basic Medicine, Shanghai Medical College, Fudan University

King Fahad Medical City

Klinisch Laboratorium ZNA

Korea Centers for Disease Control & Prevention (KCDC) Center for Laboratory Control of Infectious Diseases Division of Viral Diseases

Koupenova Lab

KRISP, KZn Research Innovation and Sequencing Platform

KRISP, KZN Research Innovation and Sequencing Platform

Kruglyak Lab

KU Leuven, Clincal and Epidemiological Virology

KU Leuven, Clinical and Epidemiological Virology

KU Leuven, Rega Institute, Clinical and Epidemiological Virology

L'institut de Recherche en Santé, de Surveillance Épidémiologique et de Formation (IRESSEF)

Lab voor klinische biologie

Lab. Microbiologia e Virologia, Cotugno, A.O. dei Colli

LABBE, Federal University of Pernambuco

LABCOVID_HCPA

Labo Klinische Biologie, UZA

Laboratoire de biologie moléculaire - Plateforme Clinique

Laboratoire de Biotechnologie

Laboratoire de microbiologie CHU NIMES

Laboratoire de Recherche et d'Analyse Médicale de la Gendarmerie Royale

Laboratoire de Recherche et d'Analyses Medicales de la Gendarmerie Royale

Laboratoire de Recherche et d'Analyses Médicales de la Gendarmerie Royale

Laboratoire de santé publique du Québec

Laboratoire de virologie - École Nationale Vétérinaire de Toulouse

Laboratoire des Procédés de Criblage Moléculaire et Cellulaire-Centre de Biotechnologie de Sfax

Laboratoire National de Sante, Microbiology, Epidemiology and Microbial Genomics

Laboratoire national de santé, Microbiology, Epidemiology and Microbial Genomics

Laboratoire National de Santé, Microbiology, Epidemiology and Microbial Genomics

Laboratoire national de sante, Microbiology, Microbial Genomics Platform

Laboratoire national de santé, Microbiology, Microbial Genomics Platform

Laboratoire Nationale de Santé, Microbiology, Epidemiology and Microbial Genomics

Laboratoire Sciences et Technologies de la Santé (STS) Institut Supérieur des Sciences de la Santé Université Hassan 1er, Settat, Morocco

Laboratoire Virpath, CIRI U111, UCBL1, INSERM, CNRS, ENS Lyon

Laboratori de Referencia de Catalunya

Laboratorio Aziendale di Microbiologia e Virologia, Azienda Sanitaria dell'Alto Adige

Laboratorio Biologia Molecolare Sars Cov2 - UOC Laboratorio Analisi - Servizio Medicina di Laboratorio, Ospedale "San Francesco" - ATS-ASSL Nuoro

Laboratorio Central de Epidemiología-DLVIE / Laboratorio de Secuenciación-Centro de Instrumentos. Instituto Mexicano del Seguro Social

Laboratorio Central de Salud Publica de Paraguay

Laboratorio Central Mg. Luis Alfredo Pianciola on behalf of 'Proyecto Argentino Interinstitucional de genomica de SARS-CoV-2' (PAIS Consortium)

Laboratório de Bioinformática e Biotecnologia (Labinftec/UFT)

Laboratório de Biologia Integrativa

Laboratório de Biologia Integrativa, Instituto de Ciências Biológicas, Universidade Federal de Minas Gerais

Laboratorio de Ecologia de Doencas Transmissiveis na Amazonia, Instituto Leonidas e Maria Deane - Fiocruz Amazonia

Laboratorio de Enfermedades Emergentes y Reemergentes

Laboratorio de Estudos de Virus Emergentes

Laboratorio de Estudos de Virus Emergentes - UNICAMP

Laboratório de Estudos de Vírus Emergentes - UNICAMP

Laboratorio de Genómica Microbiana, Universidad Peruana Cayetano Heredia

Laboratório de Imunofarmacologia

Laboratório de Imunofarmacologia - Instituto Oswaldo Cruz

Laboratorio de Infecciones Respiratorias Agudas

Laboratorio de Infecciones Respiratorias Agudas. Centro Nacional de Salud Publica,Instituto Nacional de Salud

Laboratorio de Infectología Molecular Departamento de Bioquímica y Medicina Molecular Facultad de Medicina - Universidad Autónoma de Nuevo León

Laboratorio de Infectología Molecular, Departamento de Bioquímica y Medicina Molecular,Facultad de Medicina - Universidad Autónoma de Nuevo León

Laboratório de Microbiologia Molecular - Universidade FEEVALE

Laboratório de Parasitologia Médica - Instituto de Medicina Tropical - Universidade de São Paulo

Laboratorio de Referencia Nacional de Biotecnología y Biología Molecular. Centro Nacional de Salud Publica. Instituto Nacional de Salud Peru.

Laboratorio de Referencia Nacional de Biotecnologia y Biologia Molecular. Instituto Nacional de Salud Peru

Laboratorio de Referencia Nacional de Biotecnología y Biología Molecular. Instituto Nacional de Salud Peru

Laboratorio de Referencia Nacional de Biotecnología y Biología Molecular. Instituto Nacional de Salud Perú

Laboratorio de Referencia Nacional de Biotecnología y Biología Molecular. Instituto Nacional de Salud Perú.

Laboratorio de Referencia Nacional de Biotecnología y Biología Molecular. Instituto Nacional de Salud.

Laboratorio de Referencia Nacional de Biotecnología y Biología Molecular. Instituto Nacional de Salud.Perú

Laboratorio de Referencia Nacional de Biotecnologia y Biologia Molecular.Instituto Nacional de Salud.Peru

Laboratorio de Referencia Nacional de Biotecnologia y Biologia Molecular.Instituto Nacional de Salud.Perú

Laboratorio de Referencia Nacional de Enteropatógenos. Instituto Nacional de Salud del Perú

Laboratorio de salud pública (Bogotá) and Gencore (Universidad de los Andes)

Laboratorio de Secuenciación-Centro de Instrumentos, IMSS México /Unidad de Investigación Médica en Inmunoquímica, UMAE Hospital de Especialidades “Bernardo Sepúlveda Gutiérrez”, Centro Médico Nacional Siglo XXI, Instituto Mexicano del Seguro Social (IMSS), México.

Laboratorio de Virología / Hospital Universitario Central de Asturias (HUCA)

Laboratorio de Virologia del HUCA

Laboratorio de Virología Hospital Universitario Central de Asturias (HUCA)

Laboratorio de Virologia HUCA

Laboratorio de Virología HUCA

Laboratorio de Virología Molecular

Laboratorio de Virología y Microbiología Molecular, Depto. de Microbiología, Facultad de Medicina, Universidad de El Salvador/INS-laboratorio de Ref. Ministerio de Salud

Laboratorio de Virología y Microbiología Molecular, Depto. de Microbiología, Facultad de Medicina, Universidad de El Salvador/INS-laboratorio de Ref. Ministerio de Salud.

Laboratorio de Virologia, Faculdade de Medicina, Universidade Federal de Mato Grosso, campus Cuiabá

Laboratorio de Virología, Hospital de Niños Ricardo Gutiérrez, CABA, Argentina.

Laboratorio di Genetica Medica Matera, Italy

Laboratorio di Genomica ed Epigenomica sistema Argo, Area SciencePark, Trieste, Italy;

Laboratorio di Microbiologia

Laboratorio di Microbiologia e Virologia, Università Vita-Salute San Raffaele, Milan

Laboratorio di Microbiologia e Virologia, Università Vita-Salute San Raffaele, Milano

Laboratorio di Riferimento Regionale della Sicilia Occidentale per l’Emergenza COVID-19

Laboratório Metabolismo Macromolecular FirminoTorres de Castro, Instituto de Biofísica Carlos Chagas Filho, Universidade Federal do Rio de Janeiro

laboratorio micorbiologia PO Cardarelli

Laboratorio Microbiologia e Virologia P.O. Cotugno A.O. dei Colli

Laboratorio Microbiologia e Virologia, P.O. Cotugno, A.O. dei Colli

Laboratorio Microbiologia P.O. Cardarelli

laboratorio microbiologia PO Cardarelli

laboratorio Microbiologia PO Cardarelli

Laboratorio microbiologia PO Cardarelli

Laboratorio Mixto de Biotecnología Acuática (LMBA)

Laboratorio Mixto de Biotecnología Acuática (LMBA) on behalf of 'Proyecto Argentino Interinstitucional de genomica de SARS-CoV-2' (PAIS Consortium)

Laboratorio Specialistico di Ematologia, Ospedale "San Francesco", via Mannironi 1, 08100 Nuoro

Laboratorio Specialistico Ematologia, UOC Ematologia, Ospedale "San Francesco" - ATS-ASSL Nuoro

LABORATORIO SPECIALISTICO UOC EMATOLOGIA- Ospedale San Francesco - ATS-ASSLNUoro

Laboratorio specialistico UOC Ematologia - Ospedale "San Francesco" - ATS-ASSL Nuoro

Laboratorio specialistico UOC Ematologia - Ospedale "San Francesco" - ATS-ASSL Nuoro Nuoro

Laboratorio Specialistico UOC Ematologia Ospedale San Francesco - ATS ASSL NUORO

Laboratorio SPOKE Biologia Molecolare -Azienda Ospedaliero Universitaria - AOU – Cagliari

Laboratórios de Genômica Funcional (FCA/UNESP) e Biologia Molecular (FMB-HC/UNESP) - Rede de Vigilância Genômica (Vigenômica)/UNESP

Laboratórios Genômica Funcional (FCA/UNESP) e Biologia Molecular (FMB-HC/UNESP) - Rede de Vigilância Genômica (Vigenômica)/UNESP

Laboratorium Kesehatan Provinsi Jawa Barat; Molecular Genetics Laboratory-Faculty of Medicine-Universitas Padjadjaran; School of Life Sciences and Technology & School of Pharmacy-Institut Teknologi Bandung

Laboratorium Kesehatan Provinsi Jawa Barat; School of Life Sciences and Technology & School of Pharmacy-Institut Teknologi Bandung; Molecular Genetics Laboratory-Faculty of Medicine-Universitas Padjadjaran

Laboratory Cell Biology of Viral Infection-INSERM unit 944

Laboratory Diagnostic

Laboratory Diagnostic, Veterinary Specialized Institute Kraljevo

Laboratory for advanced genomics

Laboratory for Clinical Immunology and Molecular Genetics -University Clinic Golnik

Laboratory for Functional Genome Analysis, Dept. Genomics, Gene Center of the LMU Munich

Laboratory for HIV and opportunistic infections diagnosis The Republican Research and Practical Center for Epidemiology and Microbiology (RRPCEM)

Laboratory Medicine, University of Washington

Laboratory of Bioinformatics and Computational Biology, A.C.Camargo Cancer Center

Laboratory of Biology, Department of Medicine, Democritus University of Thrace

Laboratory of Biology, Department of Medicine, Democritus University of Thrace, Alexandroupolis, Greece

Laboratory of Biology, Department of Medicine, Democritus University of Thrace, Greece

Laboratory of Cell Biology of viral infection, Unit INSERM-U944

Laboratory of Clinical Microbiology, Virology and Bioemergencies, ASST Fatebenefratelli Sacco - Sacco Hospital

Laboratory of Communicable Diseasea

Laboratory of Communicable Diseases

Laboratory of Genetics and Personalized Medicine, Zan Mitrev Clinic

Laboratory of Genomics & Bioinformatics, Institute of Immunology and Experimental Therapy, Polish Academy of Sciences

Laboratory of Genomics and Bioinformatics of the Forest Research Institute of the NAS of Belarus

Laboratory of Genomics and Bioinformatics, Comenius University Science Park

Laboratory of genomics and metagenomics

Laboratory of genomics and metagenomics, Institute of Microbiology, University Hospital Centre and University of Lausanne, Switzerland

Laboratory of Histology-Embryology

Laboratory of Histology-Embryology, Molecular Carcinogenesis Group, Faculty of Medicine, National and Kapodistrian University of Athens

Laboratory of Infectious Diseases Center of Beijing Ditan Hospital

Laboratory of Infectious Diseases, Department of Biomedical and Clinical Sciences L. Sacco, University of Milan

Laboratory of Medical Microbiology, University of Antwerp, Campus Drie Eiken, S6.26, Universiteitsplein 1, 2610, Wilrijk, Antwerp, Belgium

Laboratory of molecular-genetic research, National Center for Expertise, Kazakhstan National Center for Biotechnology, Kazakhstan

Laboratory of molecular-genetic research, National Center of Expertise, Kazakhstan National Center for Biotechnology, Kazakhstan

Laboratory of Molecular Biology and Cancer Immunology,Lebanese University Public Health England

Laboratory of Molecular Biology, Blood Center of Ribeirão Preto, Faculty of Medicine of Ribeirão Preto, University of São Paulo

Laboratory of Molecular Genetics, 2nd Faculty of Medicine, Charles University in Prague, Prague, Czech Republic

Laboratory of Molecular Virology, Department of Biomedical, Surgical and Dental Sciences University of Milano

Laboratory of Oncology, Blood Center of Ribeirão Preto, Ribeirão Preto School of Medicine, University of São Paulo

Laboratory of Parasitic Diseases, Systems Genomics Section, National Institute of Allergy and Infectious Diseases, Nathional Institutes of Health

Laboratory of Parasitic Diseases, Systems Genomics Section, National Institute of Allergy and Infectious Diseases, National Institutes of Health

Laboratory of Recombinant Vaccines

Laboratory of Respiratory Viruses and Measles, Oswaldo Cruz Institute, FIOCRUZ

Laboratory of Respiratory Viruses Teaching and Clinical Center of the Medical University of Lodz

Laboratory of Virology

Laboratory of virology and molecular diagnostics

Laboratory of virology and molecular diagnostics, Institute of Public Health

Laboratory of Virology and Molecular Diagnostics, Institute of Public Health of Republic of North Macedonia

Laboratory of Virology, INMI Lazzaro Spallanzani IRCCS

Laboratory Services Section, Texas Department of State Health Services

Laboratory Services Section, Texas Department of State Health Services-SARS-CoV-2 submission group

Laboratory, The Bio Arte Limited

LABRESIS_HCPA

LATE - Laboratório de Técnicas Especiais - Hospital Israelita Albert Einstein

Latvian Biomedical Research and Study Centre

Lauring Lab, University of Michigan, Department of Microbiology and Immunology

LBM ALPHABIO

LBM ALPHABIO, Marseille

Lebanese American University

Lednicky lab

Lednicky Laboratory at Emerging Pathogens Institute

Lednicky Laboratory at Emerging Pathogens Institute, University of Florida

Lednicky Laboratory, Emerging Pathogens Institute, University of Florida.

Lee Lab

LHUB-ULB

Li Ka Shing Faculty of Medicine, The University of Hong Kong

LIC

Lithuanian University of Health Sciences

Lithuanian University of Health Sciences Hospital, Department of Genetics and Molecular Medicine

Lithuanian University of Health Sciences, Laboratory of Molecular Cardiology

Lithuanian University of Health Sciences, Molecular cardiology lab.

LNCC

Los Alamos National Laboratory Bioscience Division

Los Angeles County PHL

Los Angeles County Public Health Laboratories, Los Angeles County Public Health Lab microbial pathogen submission group

Los Angeles County Public Health Laboratory

LSUHS Emerging Viral Threat Laboratory

Maine Health and Environmental Testing Laboratory

Mako Medical Laboratories

Malaria Research and Training Center (MRTC-Parasito)

Malaysia Genome Institute

Maryland Department of Health

Maryland Department of Health Laboratories Administration

Maryland Department of Health Laboratory

Maryland Genomics, Institute for Genome Sciences, University of Maryland School of Medicine

Maryland Public Health Laboratory

Maryland Public Health Laboratory (MD PHL)

Mason Lab

Massachusetts State Public Health Laboratory

Maximum Containment Laboratory, National Institute of Virology

MB1554

McMaster University

MD PHL

MD Public Health Laboratory

MDRU-DHR,Gandhi Medical College

MDU-PHL

MDU-PHL, The Peter Doherty Institute for Infection and Immunity

Medical Ain Shams Research Institute (MASRI), Ain Shams University

Medical Biotechnology Laboratory, Rabat Medical and Pharmacy School, Mohammed The Vth University in Rabat

Medical Genetics Department, Kocaeli University

Medical Genetics Laboratory, Regional Centre of Medical Genetics, Emergency County Hospital Craiova

Medical Genetics, Pamukkale University

Medical Laboratory Sciences, Arab American University

Medical Microbiology, Leiden University Medical Center

Medical Microbiology, Maastricht University Medical Centre

Medical Microbiology, Radboud University Medical Center

Medical Research Center, Faculty of Medicine, Syarif Hidayatullah State Islamic University Jakarta

Medtimes Molecular Laboratory

Melbourne Diagnostic Unit Public Health Laboratory (MDU-PHL)

MEPHI Aix Marseille University (AMU)

MEPHI, Aix Marseille University

Mert Kuskucu, Yesim Tuyji Tok, Kenan Midilli

MGZ Medical Genetics Center

Michigan Department of Health and Human Services

Michigan Department of Health and Human Services, Bureau of Laboratories

Micrbiological Laboratory,Lu`an Center for Disease Control and Prevention

Micrbiology Laboratory,Lu`an Center for Disease Control and Prevention

Microbial Genome Sequencing Center

Microbial Genome Sequencing Center, Microbial Genomic Epidemiological Laboratory

Microbial Genome Sequencing Center; Microbial Genomic Epidemiology Laboratory

Microbial Genome Sequencing Center; Microbial Genomic Epidemiology Laboratory, University of Pittsburgh

Microbial Genomic Epidemiology Laboratory, University of Pittsburgh

Microbial Genomics Core Lab, National Taiwan University Centers of Genomic and Precision Medicine

Microbial genomics lab LAU

Microbial Genomics lab LAU

Microbial Genomics Lab LAU

Microbial Genomics Lab LAU Byblos

Microbial Genomics Lab, Lebanese American University, Byblos

Microbial Genomics Laboratory

Microbial Genomics Laboratory, Institut Pasteur de Montevideo

Microbial Genomics Laboratory, Institut Pasteur de Montevideo, Montevideo, Uruguay

Microbial Genomics Laboratory, Institut Pasteur Montevideo

Microbial Genomics Laboratory, Institut Pasteur Montevideo, Uruguay

Microbial Pathogenomics Lab

Microbiologia Molecular, Instituto SELADIS, Universidad Mayor de San Andrés

Microbiological Diagnostic Unit - Public Health Laboratory (MDU-PHL)

Microbiological Diagnostic Unit - Public Health Laboratory (MDU-PHL), The Peter Doherty Institute for Infection and Immunity

Microbiological Diagnostic Unit Public Health Laboratory

Microbiological Diagnostic Unit Public Health Laboratory (MDU-PHL) and Victorian Infectious Disease Reference Laboratory (VIDRL)

Microbiological Diagnostic Unit Public Health Laboratory and Victorian Infectious Diseases Reference Laboratory, Doherty Institute

Microbiological Diagnostic Unit Public Health Laboratory and Victorian Infectious Diseases Reference Laboratory, The Peter Doherty Institute for Infection & Immunity

Microbiological Diagnostic Unit Public Health Laboratory and Victorian Infectious Diseases Reference Laboratory, The Peter Doherty Institute for Infection and Immunity

Microbiological Diagnostic Unit Public Health Laboratory, The Peter Doherty Institute for Infection and Immunity

Microbiological Laboratory,Lu`an Center for Disease Control and Prevention

Microbiology

Microbiology & Bioinformatics and Biostatistics, Kohat University of Science and Technology (Pakistan) & Shanghai Jiao Tong University (China)

Microbiology & Immunology, University of North Carolina

Microbiology and Immunology department, Pasteur institute in Ho Chi Minh city

Microbiology and Immunology, The Peter Doherty Institute for Infection and Immunity

Microbiology and Immunology, University of South Alabama

Microbiology and Infections Diseases

Microbiology and Infectious Diseases Dpt

Microbiology and Virology Laboratory, "Policlinico Riuniti, Azienda Ospedaliero Universitaria, Foggia"

Microbiology and Virology Unit, Florence Careggi University Hospital

Microbiology Department

Microbiology Department, University Hospital Donostia

Microbiology Department. Complexo Hospitalario Universitario de Vigo

Microbiology Division

Microbiology Division, SC DHEC

Microbiology Division, South Carolina Department of Health and Environmental Control

Microbiology Division, South Carolina Department of Health and Environmental Control (SC DHEC)

Microbiology Division, South Carolina Department of Health and Environmental Control Public Health Laboratory (SC DHEC PHL)

Microbiology laboratory- Sultan Qaboos University Hospital

Microbiology Laboratory,Lu`an Center for Disease Control and Prevention

Microbiology RPAH

Microbiology Service, Hospital Universitario Clinico San Cecilio, Granada

Microbiology, Canterbury Health Laboratories

Microbiology, Infectious Diseases and Immunology, Centre de Recherche du Centre Hospitalier de l'Universite de Montreal

Microbiology, Koc University

Microbiology, The University of Hong Kong

Microbiology, University Hospital Donostia

Microbiology, Virology and Biemergency Laboratory-ASST FBF Sacco

Microvida

Mikrobiologie, RARI

Ministry of Health Turkey

Minnesota Department of Health, Public Health Laboratory

Missouri State Public Health Laboratory

MOA Lab

MOH - Jaber Al-Ahmad Hospital (Innovation Research Laboratory)

Molecular & Genomic Pathology Laboratory, Thomas Jefferson University Hospital

Molecular Biology and Biotechnology Lab II

Molecular Biology and Virology lab, Faculty of Veterinary Medicine, Jordan University of Science and Technology

Molecular biology division, Institute of Clinical Biochemistry and Diagnostics, Charles University, Faculty of Medicine in Hradec Králové and University Hospital Hradec Králové

Molecular Biology Laboratory

Molecular Biology, IZS Sicilia

Molecular diagnostic unit for viral haemorrhagic fevers and emerging viruses, Bouaké CHU Laboratory

Molecular Diagnostics Department, Central public health laboratory

Molecular Genetics

Molecular Genetics Lab

Molecular Genetics Laboratory-Faculty of Medicine-Universitas Padjadjaran; School of Life Sciences and Technology & School of Pharmacy-Institut Teknologi Bandung; Laboratorium Kesehatan Provinsi Jawa Barat

Molecular Genetics Laboratory, Instituto de Investigaciones Químicas, Universidad Mayor de San Andrés

Molecular Infectious Disease

Molecular Microbiology & Immunology, University of Missouri

Molecular Microbiology Laboratory

Molecular Pathology Division, Department of Pathology, Hong Kong Sanatorium & Hospital

Molecular Pathology, Pathology and Laboratory Medicine Institute, Cleveland Clinic, Ohio, USA

Molecular Surveillance lab Sheikh Khalifa Medical City

Molecular Virology Laboratory of Oswaldo Cruz Foundation of Rondônia

Molecular Virology Unit, Microbiology and Virology Depertment, Fondazione IRCCS Policlinico San Matteo, Pavia

Molecular, ProgenaBiome

Molecular/Surveillance lab Sheikh Khalifa Medical City

Montana Public Health Laboratory

Monterey County Public Health Laboratory

MRC-University of Glasgow Centre for Virus Research

MRC/UVRI & LSHTM Uganda Research Unit

MRCG at LSHTM Genomics lab

MSHS Pathogen Surveillance Program

Multidisciplinary Research Unit, DHR-ICMR, Institute of Medical Sciences, Banaras Hindu University

Multidisciplinary Research Unit, DHR-ICMR, Institute of Medical Sciences, Banaras Hindu University, Varanasi

Multidrug-Resistant Organism Repository and Surveillance Network (MRSN)

Münch Lab / Ulm University Medical Center Kirchhoff Lab / Ulm University Medical Center Sparrer Lab / Ulm University Medical Center Blum Lab / LMU Munich

MUSC Molecular Pathology Laboratory

MVZ Laborärzte Singen

National Center for Expertise, Kazakhstan National Center for Biotechnology, Kazakhstan

National Center for Expertise, Kazakhstan Zonal Virology Laboratory

National Center for Expertise, National Center for Biotechnology, Kazakhstan

National Center for Infectious and Parasitic Diseases

National Center for Infectious and Parasitic Diseases (NCIPD)

National Center of Infectious and Parasitic Diseases

National Centre for Biological Sciences

National Centre For Cell Science

National Centre For Cell Science – INSACOG

National Centre for Communicable Deseases (NCCD)

National Centre for Communicable Disease (NCCD) National Influenza Center

National Centre for Communication Disease (NCCD) National Influenza Center

National Centre for Foreign Animal Disease, Canadian Food Inspection Agency

National Centre for Infectious Diseases, National Centre for Infectious Diseases

National Genomics Core- Center for DNA Fingerprinting and Diagnostics (NGC-CDFD)- DBT's PAN-INDIA-1000 Genome consortium

National Health Laboratory Service (NHLS), Tygerberg

National Health Laboratory Service/UCT

National Health Laboratory Services

National Health Laboratory Services, Virology

National Influenza Center

National Influenza Center - National Institute of Hygiene and Epidemiology (NIHE)

National Influenza Center, Bahrain

National Influenza Center, Indian Council of Medical Research-National Institute of Virology

National Influenza Center, Indian Council of Medical Research - National Institute of Virology

National Influenza Center, National Institute of Hygiene and Epidemiology (NIHE)

National Influenza Centre for Northern Greece

National Influenza Centre for Nothern Greece

National Institute for Allergy and Infectious Diseases Integrated Research Facility - Frederick (NIAID IRF- Frederick), National Institutes of Health (NIH)

National Institute for Communicable Disease Control and Prevention (ICDC) Chinese Center for Disease Control and Prevention (China CDC)

National Institute for Communicable Disease Control and Prevention, Chinese Center for Disease Control and Prevention

National Institute for Communicable Diseases of the National Health Laboratory Service

National Institute for Infectious Diseases, INMI, "L. Spallanzani" IRCCS

National Institute for Public Health and the Environment (RIVM)

National Institute for Viral Disease Control & Prevention, CCDC

National Institute for Viral Disease Control & Prevention, China CDC

National Institute for Viral Disease Control and Prevention

National Institute for Viral Disease Control and Prevention, China CDC

National Institute for Viral Disease Control and Prevention, China CDC, Yunnan Provincial CDC

National Institute of Biomedical Genomics

National Institute of Biomedical Genomics - DBT's PAN-INDIA 1000 SARS--CoV-2 RNA Genome Sequencing Consortium

National Institute of Biomedical Genomics – INSACOG

National Institute of Biotechnology

National Institute of Health Research and Development

National Institute of Health, Department of Medical Sciences, Ministry of Public Health, Thailand

National Institute of Health. Department of medical Sciences, Ministry of Public Health, Thailand

National Institute of Hygiene and Epidemiology (NIHE)

National Institute of Infectious Diseases-Prof. Dr. Matei Bals Molecular Diagnostics Laboratory

National Institute of Infectious Diseases (NIID)

National Institute of Public Health

National Institute of Public Health - National Institute of Hygiene

National Institute of Public Health - National Institute of Hygiene, DNA Sequencing and Synthesis Facility (oligo.pl) - Institute of Biochemistry and Biophysics PAS

National Institute of Virology-Microbial Containment Complex, Indian Council of Medical Research

National Institute of Virology, NIV Influenza

National Institute of Virology, Pune

National Key Laboratory of Gene Technology, Institute of Biotechnology (IBT)

National Key Laboratory of Gene Technology, Institute of Biotechnology, Vietnam Academy of Science and Technology

National Laboratory for Health, Environment and Food

National Laboratory for Health, Environment and Food;Institute of Microbiology and Immunology

National Laboratory of Virology, Szentágothai Research Centre

National Medical Research Center for Obstetrics, Gynecology and Perinatology named after Academician V.I.Kulakov of the Ministry of Healthcare of Russian Federation

National Microbiology Laboratory

National Microbiology Laboratory (NML)

National Plateform bis UMONS/Jolimont

National Platform bis UMONS/Jolimont

National Public Health Center, National Biosafety Laboratory

National Public Health Laboratory

National Public Health Laboratory, National Centre for Infectious Diseases

National Public Health Organization

National Public Health Surveillance Laboratory

National Reference Center for Viruses of Respiratory Infections, Institut Pasteur, Paris

National Reference Laboratory

National Reference Laboratory for COVID-19, Pasteur Institute of Iran

National Reference Laboratory for Influenza and Respiratory Viruses CZE

National Reference Laboratory, NCDC, Abuja

National reference Laboratory, NCDC, Gaduwa, Abuja

National Reference Laboratory, NCDC, Gaduwa, Abuja

National reference Laboratory, NCDC, Gaduwa, Abuja, Nigeria

National Reference Laboratory, NCDC, Gaduwa, Abuja, Nigeria

National Reference Laboratory, Nigeria Centre for Disease Control, Gaduwa, Abuja, Nigeria

National Reference Laboratory, Nigeria Centre for Disease Control, Gaduwa,Abuja, Nigeria

National Research Center for Translational Medicine (Shanghai), Ruijin Hospital affiliated to Shanghai Jiao Tong University School of Medicine & Shanghai Public Health Clinical Center

National Veterinary Institute

National Virology Reference Laboratory

National Virus Reference Laboratory

Naval Medical Research Center Biological Defense Research Directorate

NBACC

NBCC Sequencing Facility

NCDC Institute of Genomics and Integrative Biology

NCDC/CSIR-IGIB

NCDC/IGIB

NCSLPH

Nebraska Public Health Laboratory COVID-19 Response Team

Nepal Health Research Council

Nevada State Health Lab

Nevada State Public Health Laboratory

New Jersey Public Health and Environmental Laboratories

New Jersey Public Health and Environmental Laboratories (NJ PHEL)

New Jersey Public Health and Environmental Laboratories (NJPHEL)

New Jersey Public Health and Environmental Laboratories (PHEL)

New Jersey Public Health Environmental Laboratories (NJ_PHEL)

New Mexico Department of Health Scientific Laboratory

New York City Public Health Laboratory

New York Genome Center

Next Generation Sequencing Reference Laboratory, Faculty of Medicine, Cairo University and The Center for Genome and Microbiome Research, Faculty of Pharmacy, CAIRO UNIVERSITY

Next Generation Sequencing Reference Laboratory, Faculty of Medicine, CAIRO UNIVERSITY and The Center for Genome and Microbiome Research, Faculty of Pharmacy, CAIRO UNIVERSITY

NGS Competence Center Tübingen, Institut für Medizinische Mikrobiologie und Hygiene, Universitätsklinikum Tübingen

NGS Competence Center Tuebingen, Institut für Medizinische Mikrobiologie und Hygiene, Universitaetsklinikum Tübingen

NGS Lab, DNA SOLUTION LTD.

NHLS/UCT

Nigerian Institute of Medical Research

NIV Influenza

NJ Public Health and Environmental Laboratories

NJ_PHEL

NLZOH (National Laboratory for Health, Environment and Food) / CISLD (Clinical Institute of Special Laboratory Diagnostics), University Children's Hospital, University Medical Center Ljubljana

NLZOH, Laboratory for Virology

No.409,Gaocheng Middle Road,Lu`an City,Anhui Provice

Nodo de Secuenciación Tierra del Fuego - Hospital Regional Ushuaia - Centro Austral De Investigaciones Científicas - Universidad Nacional De Tierra Del Fuego on behalf of 'Proyecto Argentino Interinstitucional de genomica de SARS-CoV-2' (PAIS Consortium)

North Carolina State Laboratory of Public Health

North Dakota Department of Health, Public Health Laboratory

Northumbria University

Northwestern University - Ozer Lab

Norwegian Institute of Public Health

Norwegian Institute of Public Health, Department of Virology

Notre Dame Genomics & Bioinformatics Core Facility

NPHL

NPHL COVID-19 Response Team

NRL-HIV

NSU Genome Research Institute (NGRI)

NSU Genome Research Institute (NGRI), North South University

NSW Health Pathology - Institute of Clinical Pathology and Medical Research; Centre for Infectious Diseases and Microbiology Laboratory Services; Westmead Hospital; University of Sydney

NSW Health Pathology - Institute of Clinical Pathology and Medical Research; Westmead Hospital; University of Sydney

Ohio Depart of Health Laboratory

Ohio Department of Health

Ohio Department of Health Laboratories

Ohio Department of Health laboratory

Ohio Department of Health Laboratory

Oklahoma Animal Disease Diagnostic Laboratory

Oklahoma Animal Disease Diagnostic Laboratory, Oklahoma State University

OLVZ Aalst

Oman-National Influenza Center

Oman-NIC

Omics Sciences Lab

Omics Sciences Laborator

Omics Sciences Laboratory

ONCOGENE LLC

Onderzoeksgroep Virologie

Ontario Agency for Health Protection and Promotion (OAHPP)

Ontario Institute for Cancer Research

Orebro University Hospital

Oregon SARS-CoV-2 Genome Sequencing Center

Oregon State Public Health Laboratory

Osmania University

OSU Center for Genome Research and Biocomputing

OSU Polaris Molecular Laboratory

OUCRU

OUCRU/HTD

Oxford University Clinical Research Unit (OUCRU)

Oxford University Clinical Research Unit, Hanoi, Vietnam

Ozer Lab

Pamukkale University Department of Medical Genetics

Pandemic Response Lab, R&D

Pathogen Discovery

Pathogen Discovery, Respiratory Viruses Branch, Division of Viral Diseases, Centers for Disease Control and Prevention

Pathogen Genomics Center, National Institute of Infectious Diseases

Pathogen Genomics Lab King Abdullah University of Science and Technology (KAUST)

Pathogen Genomics Lab King Abdullah University of Science and Technology(KAUST)

Pathogen Laboratory (BSL3), Biomedical Innovation Department, Experimental and Applied Biology Division, Scientific Research Center and High Education from Ensenada (CICESE)

Pathogen Sequencing Lab, National Institute for Biomedical Research (INRB)

Pathogenic Microorganisms Variability Laboratory

Pathology and Laboratory Medicine Institute, Cleveland Clinic, Ohio, USA

Pathology and Laboratory Medicine, UW-Madison

PathWest Laboratory Medicine WA

PathWest Laboratory Medicine WA Microbial Surveillance Unit

PCR Laboratory, The First Affiliated Hospital of Zhengzhou University, Zhengzhou, Henan, China

Philippine Genome Center

Philippine Genome Center, University of the Philippines System

PHV-FSS

Physiology

Physiology, Istanbul Medeniyet University

Piantadosi Lab, Emory Department of Pathology

Pirogov Russian National Research Medical University, Research and Development

Planet Lab, Children's Hospital of Philadelphia

Plateforme Clinique de testing Namuroise

Plateforme de testing Namuroise

Polaris Molecular Laboratory

Population Medicine and Diagnostic Sciences, Cornell University

Princess Haya Biotechnology Center, Jordan University of Science and Technology

Princess Haya Biotechnology Center/ Jordan University of Science & Technology

Pro-Vitam Diagnostics and Research Laboratory

Prof. Gorgoulis Lab

Prof. Massimo Zollo CEINGE TASK-FORCE COVID19 - Regione Campania

Programa de Oncovirologia, Instituto Nacional de Câncer

Programme in Emerging Infectious Diseases, Duke-NUS Medical School

Project group Epidemiology of Highly Pathogenic Microorganisms, Robert Koch-Institute

Project group Epidemology of Highly Pathogenic Microorganisms, Robert Koch-Institute

Project group Epidemology of Highly Pathogenic Microorganisms, Robert Koch Institut

Protzer Lab

Protzer Lab, Gagneur Lab, Robert Koch Institut

Protzer Lab, Institut für Medizinische Mikrobiologie und Hygiene, Gagneur Lab

Protzer Lab, Laboratory for Functional Genome Analysis, Dept. Genomics, Gene Center of the LMU Munich

Providence St. Joseph Health Molecular Genomics Laboratory

Public Health Agency of Canada - National Microbiology Laboratory

Public Health Lab

Public Health Laboratory - Infectious Disease Lab, Minnesota Department of Health Infectious Disease Laboratory Submission Group

Public Health Laboratory, Saudi CDC

Public Health Ontario Laboratories

Public Health Ontario Laboratory

Public Health Virology-Forensic and Scientific Services

Public Health Virology-Forensic and Scientific Services (PHV-FSS)

Public Health Virology Laboratory

Public Health Virology Laboratory, Forensic and Scientific Services (PHV-FSS)

Public Health Virology Laboratory, Forensic and Scientific Services, Queensland Health

Public Health Virology Laboratory, Forensics and Scientific Services, Queensland Health

Public Health Wales Microbiology Cardiff

Public Health Wales Microbiology Cardiff Wales Specialist Virology Centre

Public Health, United States Air Force School of Aerospace Medicine

Q Squared Solutions - QRTP facility

Quadram Institute Bioscience

QUALITY CONTROL CHEMICAL BIOLOGICAL RISK, AOOR Villa Sofia Cervello Palermo

Quantigen Biosciences

Queen's Genomics Lab at Ongwanada (Q-GLO)

Quest Diagnostics

R. G. Lugar Center for Public Health Research, National Center for Disease Control and Public Health (NCDC) of Georgia.

Radboudumc

Rafik Hariri University Hospital

Rapid Response Team

Razi Vaccine and Serum Research Institute

RCMI-Center for Research Resources, Ponce Research Institute

Redeemer's University, ACEGID

Reditus Laboratories

Research and Experiment Center, Meizhou People Hospital

Research and Medical Analysis Laboratory of Gendarmerie Royale

Research Center for Genetic Engineering and Biotechnology "Georgi D. Efr

Research Center for Genetic Engineering and Biotechnology "Georgi D. Efremov” , Macedon

Research Center for Genetic Engineering and Biotechnology "Georgi D. Efremov” , Macedoni

Research Center for Genetic Engineering and Biotechnology "Georgi D. Efremov” , Macedonian Academ

Research Center for Genetic Engineering and Biotechnology "Georgi D. Efremov” , Macedonian Academy of Sciences and Arts

Research Center for Vaccine Technology and Development, Institute of Tropical Disease, Airlangga University

Research Institute for Tropical Medicine

Research platform for Transfusion-transmitted Disease, Institute of Blood Transfusion, Chinese Academy of Medical Sciences

Research Unit of Systems Microbiology

Respiratory virus Laboratory, Chinese Academy of Medical Science

Respiratory Virus Unit, Microbiology Services Colindale, Public Health England

Respiratory Virus Unit, National Infection Service, Public Health England

Respiratory Viruses Branch, Centers for Disease Control and Prevention

Respiratory Viruses Branch, Division of Viral Diseases, Centers for Disease Control and Prevention

Riga East University Hospital-National Microbiology Reference Laboratory; Eurofins Genomics Europe Sequencing GmbH

RIPHL at Rush University Medical Center

Robert Koch Institut

Robert Koch Institute

Robert Koch Institute, Bioinformatics MF1, Berlin, Germany

Robert Koch Institute, Influenza and respiratory viruses FG17 & Bioinformatics MF1, Berlin, Germany

Robert Koch Institute, ZBS1 Highly Pathogenic Viruses & Bioinformatics MF1, Berlin, Germany

Robert Koch Institute, ZBS1 Highly Pathogenic Viruses, Berlin, Germany

Rocky Mountain Laboratories

Rocky Mountain Laboratories, RTS Genomics Unit, National Institute of Allergy and Infectious Diseases, National Institutes of Health

Roy J. Carver Biotechnology Center

Royal Hobart Hospital

RSE "National Center for Biotechnology"

RSE "National Center for Biotechnology" and RSE "National Center of Expertise"

RSE "National Center of Expertise" and RSE "National center for Biotechnology"

Ruder Boškovic Institute; Forensic Science Centre Ivan Vucetic; University of Zagreb Faculty of Science

Rwanda National Laboratory

Rwanda National Reference Laboratory

Ryota Kumagai Tokyo Metropolitan Institute of Public Health

S.M.S. Medical College

S.M.S.Medical College, Jaipur, Rajasthan

S.S. Genetica e Tecniche Omiche Avanzate Istituto Zooprofilattico Sperimentale del Piemonte, Liguria e Valle d'Aosta

SA Pathology

Saitama Medical University

Salzkammergutklinikum Vöcklabruck, Institut für Pathologie

San Gallicano Dermatological Institute I.F.O.

Santa Clara County public Health Laboratory

Santa Clara County Public Health Laboratory

Sapporo City Institute of Public Health

SARS-CoV-2 Sequencing Castilla y Leon-Spain Consortium

SC Department of Health and Environmental Control

SC Microbiologia e Virologia AOUSS

School of Life Sciences and Technology & School of Pharmacy-Institut Teknologi Bandung; Molecular Genetics Laboratory-Faculty of Medicine-Universitas Padjadjaran; Laboratorium Kesehatan Provinsi Jawa Barat

School of Pharmacy

School of Pharmacy & School of Life Sciences and Technology - Institut Teknologi Bandung; Molecular Genetics Laboratory-Faculty of Medicine-Universitas Padjadjaran; Laboratorium Kesehatan Provinsi Jawa Barat

School of Pharmacy, Shenandoah University

School of Public Health, The University of Hon g Kong

School of Public Health, The University of Hong Kong

School of Veterinary Medicine, Disease Control

Schwessinger Lab

Scientific Veterinary Institute "Novi Sad"

SCSFRI, South China Sea Fisheries Research Institute, Chinese Academy of Fishery Sciences (SCSFRI, CAFS)

SEA Microbiome Unit, Faculty of Industrial Sciences & Technology, Universiti Malaysia Pahang

Seattle Flu Study

Seattle Flu Study, University of Washington Medical Center

Second Hospital of Anhui Medical University

Second Military Medical University, Department of Microbiology

Section for Molecular Diagnostics

SeqCOVID-SPAIN consortium / IBV (CSIC)

SeqCOVID-SPAIN consortium/IBV (CSIC)

SeqCOVID-SPAIN consortium/IBV(CSIC)

SeqCOVID-SPAIN consortium/Institute of Biomedicine of Valencia, IBV-CSIC

Sequencing and Bioinformatics Service and Molecular Epidemiology Research Group. FISABIO-Public Health

Sequencing and Bioinformatics Service and Molecular Epidemiology Research Group. FISABIO-Public Health, and SeqCOVID-Spain Consortium

Sequencing and Bioinformatics Service and Molecular Epidemiology Research Group. FISABIO-Public Health.

Sequencing and Bioinformatics Service FISABIO-Public Health

Sequencing and Bioinformatics Service. Molecular Epidemiology Laboratory. FISABIO-Public Health

Servicio de Microbiología Hospital Ramón y Cajal

Servizo de Microbioloxía. Complexo Hospitalario de Santiago de Compostela

Servizo de Microbioloxía. Complexo Hospitalario de Santiago de Compostela.

Servizo de Microbioloxía. Complexo Hospitalario de Santiago de Compostela21

Shenzhen Key Laboratory of Pathogen and Immunity, National Clinical Research Center for Infectious Disease, Shenzhen Third People's Hospital

Shenzhen Key Laboratory of Pathogen and Immunity, National Clinical Research Center for Infectious Disease,Shenzhen Third People's Hospital

Sher-e-Bangla Nagar, Agargaon, Dhaka-1207, Bangladesh.

Shiraz University

Sídlištní 136/24 165 03, Prague Czech Republic

siParadigm LLC

Smith Laboratory, Centre de Recherche CHU Sainte-Justine

SMS Medical College jaipur

SMS Medical College Jaipur

SMU Metagenomics lab

SON ESPASES UNIVERSITARY HOSPITAL

South Carolina Department of Health and Environmental Control

South China Agricultural University

South Dakota Public Health Laboratory

South Dakota Public Health Laboratory, South Dakota Department of Health

Southern Nevada Public Health Laboratory

Special Infectious Agents Unit

Special Operations Medical Research Division, Defence Services Medical Research Centre

Specialized Lab for COVID-19 Detection, Department of Genetic Engineering and Biotechnology

St. Jude Children's Research Hospital Infectious Diseases

St.Vincent's University Hospital

Stanford University School of Medicine, Clinical Virology Laboratory

State Center for Health Surveillance of the Health Department of the State of Rio Grande do Sul (CEVS/SES-RS)

State Hygienic Laboratory at the University of Iowa

State Key Laboratory for Diagnosis and Treatment of Infectious Diseases, National Clinical Research Center for Infectious Diseases, First Affiliated Hospital, Zhejiang University School of Medicine, Hangzhou, China 310003

State Key Laboratory for Diagnosis and Treatment of Infectious Diseases, National Clinical Research Center for Infectious Diseases, First Affiliated Hospital, Zhejiang University School of Medicine, Hangzhou, China. 310003

State Key Laboratory for Emerging Infectious Diseases Department of Microbiology Li Ka Shing Faculty of Medicine The University of Hong Kong

State Key Laboratory of Agriculture Microbiology, Huazhong Agric

State Key Laboratory of Agriculture Microbiology, Huazhong Agric Laboratory of Animal Virology, College of Veterinary Medicine

State Key Laboratory of Biotherapy of Sichuan University

State Key Laboratory of Emerging Infectious Diseases, The University of Hong Kong

State Key Laboratory of Genetic Resources and Evolution, Kunming Institute of Zoology, Chinese Academy of Sciences

State Key Laboratory of Respiratory Disease, National Clinical Research Center for Respiratory Disease, Guangzhou Institute of Respiratory Health, the First Affiliated Hospital of Guangzhou Medical University

State Key Laboratory of Virology, Wuhan University

State Laboratories Division, Hawaii State Department of Health

State Research Center of Virology and Biotechnology VECTOR, Department of Collection of Microorganisms

State Veterinary Institute Prague

State Veterinary Institute Prague and The National Institute of Public Health

State Virus Research and Diagnostic Laboratory (VRDL), AIIMS Raipur

Statens Serum Institute

Stefan cel Mare, University Metagenomics lab

Steininger Lab

Stellenbosch University and NHLS

Stem Cell Lab, Universitas Pembangunan Nasional Veteran Jakarta

Stem Cell Lab, Universitas Pembangunan Nasional Veteran Jakarta (UPNVJ)

Stem Cell Lab. Universitas Pembangunan Nasional Veteran Jakarta (UPNVJ)

Stern Lab

Submitting lab: Laboratorio SPOKE Biologia Molecolare -Azienda Ospedaliero Universitaria - AOU – Cagliari

Swiss National Reference Centre for Influenza

Swiss National Reference Centre for Influenza Virology laboratory, CNRI

Swiss Tropical and Public Health Institute

Switzerland

Synergy Laboratories

Synergy Laboratories, Inc.

Taiwan Centers for Disease Control

Takayuki Hishiki Kanagawa Prefectural Institute of Public Health

Tampa General Hospital Esoteric Research & Development Lab

Tanjungpura University Hospital

TaskForce Covid19-Ceinge Regione Campania

Technical Support Units for Scientific Research (UATRS), National Centre for Scientific and Technical Research (CNRST)

Technology Centre, Guangzhou Customs

Tehran University of Medical Sciences

Tejgaon College bmb lab

Telethon Institute of Genetics and Medicine - Telethon Institute of Genetics and Medicine - TIGEM

Telethon Institute of Genetics and Medicine - TIGEM

Telethon Institute of Genetics and Medicine (TIGEM)

Tewhey Lab, The Jackson Laboratory

Texas A&M Institute for Genomic Sciences and Society (TIGSS)

Texas Children's Microbiome Center

Texas Department of State Health Services

Texas Department of State Health Services - TXDSHS

Texas Department of State Health Services (TXDSHS)

TGen North

Thai National Influenza Center, Department of medical Science, Ministry of Public Health, Thailand

Thai Red Cross Emerging Infectious Diseases Center and Faculty of Medicine, Chulalongkorn University

THE AFRICA GENOMICS CENTRE AND CONSULTANCY

THE AFRICA GENOMICS CENTRE AND CONSULTANCY LIMITED

The Department of Infectious Disease Prevention and Control, Henan Provincial Center for Disease Control and Prevention

the First Affiliated Hospital of Guangzhou Medical University & BGI-Shenzhen

The First Affiliated Hospital of Guangzhou Medical University & BGI-Shenzhen

The Foundation for Medical Research

The Hospital for Sick Children

The Institute of Molecular Biology and Genetics of NASU

The Jackson Laboratory

The National Institute of Public Health and State Veterinary Institute Prague

The National Institute of Public Health Center for Epidemiology and Microbiology

The National Laboratory of Health, Environment and Food - Centre for Medical Microbiology Maribor

The National Laboratory of Health, Environment and Food, Maribor, Slovenia

The Ohio State University-James Molecular Lab at Polaris

The Ohio State University Applied Microbiology Services Laboratory

The Ohio State University James Molecular lab

The Public Health Agency of Sweden

The University of Hong Kong

The University of Hong Kong Department of Microbiology

THSTI Bioassay laboratory

TIGEM

TIGSS

Tilia Laboratories s.r.o.

Tokyo Metoropolitan Institute of Public Health

Tokyo Metropolitan Institute of Public Health

Tokyo Metropolitan Institute of Public Health, Department of Microbiology

Transcriptomics & Applied Genomics (TAG)

TransVIHMI(Recherches Translationnelles sur le VIH et les Maladies Infectieuses)

TransVIHMI, IRD/INSERM/Monpellier University

TSGH-CP molecular lab

TSGH-CP molecular lab, Division of Clinical Pathology, Department of Pathology

Tumor Immunology Unit, Department of Health Sciences, University of Palermo School of Medicine; Section of Microbiology, University of Palermo School of Medicine; National Research Council of Italy - High Performance Computing and Networking Institute (CNR-ICAR) of Palermo

TwinStrand Biosciences, Inc.

TXDSHS

TxGen

U.O. Diagnostica Virologica Dip. Sanità Animale IZSM

U.O. Genomics, S.S. Genetics and Advanced Omics Techniques Istituto Zooprofilattico Sperimentale del Piemonte, Liguria e Valle d'Aosta

U.O. Genomics, S.S. Genetics and Advanced Omics Techniques, Istituto Zooprofilattico Sperimentale del Piemonte Liguria e Valle d'Aosta

U.O. Genomics, S.S. Genetics and Advanced Omics Techniques, Istituto Zooprofilattico Sperimentale del Piemonte, Liguria e Valle d'Aosta

U.O. Igiene, Ospedale Policlinico San Martino

U.O. Microbiologia Laboratorio Unico Centro Servizi AUSL della Romagna

U.O. Microbiologia, Laboratorio Unico Centro Servizi - AUSL della Romagna

U.S. Air Force School of Aerospace Medicine

U.S. Naval Medical Research Center Biological Defense Research Directorate

UAntwerp, Laboratory of Medical Microbiology

UAntwerp, Laboratory of Medical Microbiology,

UAntwerp, Laboratory of Medical Microbiology, Campus Drie Eiken S6.26, Universiteitsplein 1, 2610, Wilrijk, Antwerp, Belgium

UAntwerp, Laboratory of Medical Microbiology, Campus Drie Eiken S6.26, Universiteitsplein 1, 2610, Wilrijk, Belgium

UC Davis Genome Center

UCD National Virus Reference Laboratory

UCLouvain/IREC/MBLG

UCSC Genomics Institute

UFS Virology

UHAS COVID-19 Lab

Uhlemann Laboratory, Columbia University Irving Medical Center

UHTL

UMC Groningen, Clinical Virology, Department of Medical Microbiology and Infection Prevention

UMR 190 - Faculte de medecine, UMR 'Emergence des Pathologies Virales' (EPV: Aix-Marseille University - IRD 190 - Inserm 1207 - EHE

UMR 8199/1283 EGID

UMR PIMIT

UMR PIMIT Université de La Réunion

UMR190-Unité des virus emergents

Unidad de Genomica Avanzada

Unidad Universitaria de Secuenciación Masiva y Bioinformática (UUSMB). IBT-UNAM

Unit for Biological Agents, Department for CBRN Defence and Security, Swedish Defence Research Agency

Unit for Laboratory Development and Technology Transfer, Public Health Agency of Sweden

Unità di Analisi del Rischio ed Epidemiologia Genomica, Istituto Zooprofilattico Sperimentale dell'Emilia Romagna e della Lombardia (IZSLER)

Unite des virus emergents, UMR190

Unité Mixte Internationale TransVIHMI (UMI 233 IRD – U1175 INSERM - Université de Montpellier) IRD (Institut de recherche pour le développement)

Unité Mixte Internationale TransVIHMI (UMI 233 IRD – U1175 INSERM - Université de Montpellier)IRD (Institut de recherche pour le développement)

United States Air Force School of Aerospace Medicine

Universidad del Valle, Universidad Nacional de Colombia-Sede Palmira, International Center for Tropical Agriculture

Universidad Industrial de Santander

Universidad Industrial de Santander.

Universidad Nacional de Colombia - Laboratorio Genómico One Health

Universidade Federal de Ciências da Saúde de Porto Alegre

Universidade Federal do Parana (UFPR)

Universitas Sebelas Maret (UNS), Rumah sakit Universitas Sebelas Maret (RS-UNS) Surakarta, National Institute of Health Research and Development, Indonesian Ministry of Health

Universitas Sebelas Maret (UNS); Rumah Sakit Universitas Sebelas Maret (RS-UNS), Surakarta; National Institute of Health Research and Development, Indonesian Ministry of Health , Jakarta).

Universitas Sebelas Maret (UNS); Rumah Sakit Universitas Sebelas Maret (RS-UNS), Surakarta; National Institute of Health Research and Development, Indonesian Ministry of Health, Jakarta.

Universitas Sebelas Maret (UNS); Rumah Sakit UNS (RS-UNS); National Institute of Health Research and Development, Indonesian Ministry of Health.

Universitätstr.1 40225 Düsseldorf Germany

University at Buffalo Genomics and Bioinformatics Core

University Clinical Research Center, University of Sciences

University Hospital Basel, Clinical Bacteriology

University Hospital Basel, Clinical Virology

University Hospital Basel, Labormedizin

University Hospital Regensburg

University Hospitals of Geneva Laboratory of Virology

University Hospitals of Geneva, Laboratory of Virology

University Hospitals Translational Laboratory (UHTL), University Hospitals

University Medical Center Hamburg-Eppendorf

University of Alabama at Birmingham

University of Bari Biomedical Sciences and Human Oncology

University of California, San Francisco

University of Florida

University of Florida, Lednicky Lab

University of Iowa, McCray Lab

University of Miami Immunology and Histocompatibility Laboratory

University of Minnesota Genomics Center

University of Mississippi Medical Center, Molecular and Genomics Core Facility

University of Oregon Genomics and Cell Characterization Core Facility (GC3F)

University of Otago

University of Portsmouth

University of Sarajevo Veterinary Faculty

University of Sarajevo, Veterinary Faculty

University of Sarajevo, Veterinary Faculty, Laboratory for Molecular Diagnostic and Research Laboratory

University of South Carolina Functional Genomics Core

University of Tabriz

University of Texas at Austin Genome Sequencing and Analysis Facility (UTGSAF)

University of Texas, Genomic Sequencing and Analysis Facility

University of Ulsan College of Medicine and Asan Medical Center

University of Verona, Department of Biotechnology

University of Warwick, for the COVID-19 Genomics (COG) UK Consortium

University of Washington Virology Lab

University of Washington, Laboratory Medicine

University of Wisconsin-Madison AIDS Vaccine Research Laboratories

University of Wisconsin Madison, AIDS Vaccine Research Laboratories

University of Zagreb, Centre for research and knowledge transfer in biotechnology

University of Zambia, School of Veterinary Medicine, Disease Control

University or Iowa, Lung Biology and Cystic Fibrosis Research Center, Pezzulo Lab

UNMC COVID-19 Response Team

UNS/RS-UNS, Faculty of Medicine

UNZAVET and PATH

US Air Force School of Aerospace Medicine

Utah Public Health Laboratory

Utah Public Health Laboratory, Utah Public Health Laboratory Infectious Disease submission group

UTGSAF

UW Virology lab

UW Virology Lab

UZA, Clinical Biology

van Bakel Laboratory, Genetics and Genomics Sciences, Icahn School of Medicine at Mount Sinai

Vanda Pharmaceuticals

VETAL Animal Health Products Company, BSL3+ Production Laboratuary /Turkey

VETAL Animal Health Products Company, BSL3+ Production Laboratuary, Turkey

Veterinary Specialized Institue Kraljevo

Veterinary Specialized Institute "Kraljevo", Serbia

Veterinary Specialized Institute "Nis", Serbia

Veterinary Specialized Institute "Sabac", Serbia

Veterinary Specialized Institute Kraljevo

Victorian Infectious Diseases Reference Laboratory

Victorian Infectious Diseases Reference Laboratory (VIDRL) and the Melbourne Diagnostic Unit Public Health Laboratory (MDU-PHL)

Victorian Infectious Diseases Reference Laboratory and Microbiological Diagnostic Unit Public Health Laboratory, Doherty Institute

VIDRL and MDU-PHL

ViFU

Vilnius University Hospital Santaros Klinikos

Vilnius university hospital Santaros Klinikos, Center of Laboratory Medicine

Vilnius University Hospital Santaros Klinikos, Center of Laboratory Medicine

Viollier AG

Viral vaccines, VSVRI- Veterinary serum and vaccine research institute

Virginia DCLS

Virginia Division of Consolidated Laboratory Services

Virginia Division of Consolidated Laboratory Services (DCLS)

ViroGenetics - BSL3 Laboratory of Virology; Human Genome Variation Research Group & Genomics Centre MCB; Bioinformatics Research Group

ViroGenetics - BSL3 Laboratory of Virology; Human Genome Variation Research Group & Genomics Centre MCB; Bioinformatics Research Group Department of Virology

Virologisches Institut, Universitätsklinikum Erlangen

Virology

Virology (microbiology), Hospital Universitario Central de Asturias

Virology and Legal Medicine Laboratories, Department of Biomedical Sciences and Public Health, University Politecnica delle Marche

Virology department Institute of microbiology and immunology Faculty of Medicine University of Belgrade

Virology Department Institute of Microbiology and Immunology Faculty of Medicine University of Belgrade

Virology Department, Institute of microbiology and immunology, Faculty of Medicine University of Blegrade

Virology Department, Royal Infirmary of Edinburgh, NHS Lothian

Virology Lab,Department of Pathology, National Cheng Kung University Hospital

VIROLOGY LABORATORY-CHU NICE

Virology Laboratory, Scientific Department, Army Medical Center

Virology Research Laboratory; Area of Virology, Serology and Virology Division (SAViD), New South Wales Health Pathology Randwick

Virology Service, Centre Pasteur of Cameroun

Virology Unit, Agrobiodiversity and Biotechnology Project, CIAT - International Center for Tropical Agriculture

Virology Unit, Institut Pasteur de Madagascar

Virology Unit, Institut Pasteur du Cambodge

Virology Unit, Institut Pasteur du Cambodge (Sequencing done by: Jessica E Manning/Jennifer A Bohl at Malaria and Vector Research Research Laboratory, National Institute of Allergy and Infectious Diseases and Vida Ahyong from Chan-Zuckerberg Biohub)

Virology, CHU Pitie Salpetriere Charles Foix

Virology, Ecole Nationale Veterinaire de Toulouse

Virology, ICAR-National Research Centre on Equines

Virology, National Health Laboratory Service, Charlotte Maxeke Johannesburg Academic Institute

Virology, National Institute for Biological Standards and Control

Virology, NHLS, CMJAH

Virology, Wageningen Bioveterinary Research

Virus Ecology Section, RML

Virus Ecology, Rocky Mountain Laboratories, National Institutes of Health

Virus Research Laboratory, Department of Zoology, Osmania University, Hyderabad, India

Virus Research Laboratory, Department of Zoology, Osmania University,Hyderabad,India

VPRL

VRDL-Gandhi Medical College

WACCBIP, University of Ghana

Wadsworth Center, New York State Department of Health

Wadsworth Center, New York State Department.of Health

WallauLab, Aggeu Magalhaes Institute

Washington State Department of Health Public Health Laboratories

Washington University in St. Louis

Weifang Center for Disease Control and Prevention & BGI-Shenzhen

Wellcome Sanger Institute for the COVID-19 Genomics UK (COG-UK) consortium

Wellcome Sanger Institute for the COVID-19 Genomics UK (COG-UK) Consortium

West African Centre for Cell Biology of Infectious Pathogens (WACCBIP), University of Ghana, Volta Road, Legon-Accra, Ghana

West Java Health Laboratory

West Java Health Laboratory; School of Life Sciences and Technology, Institut Teknologi Bandung

Where sequence data have been generated and submitted to GISAID

WHO National Influenza Centre Russian Federation

Wiedenheft lab, Montana State University

Wisconsin State Laboratory of Hygiene Communicable Disease Division

Wojewodzka Stacja Sanitarno-Epidemiologiczna w Olsztynie, Laboratorium Badan Epidemiologiczno-Klinicznych

Wojewódzka Stacja Sanitarno-Epidemiologiczna w Olsztynie, Laboratorium Badan Epidemiologiczno-Klinicznych

Worobey Lab on behalf of the Arizona COVID-19 Genomics Union

Worobey Lab, Department of Ecology and Evolutionary Biology, University of Arizona

Wuhan Institute of Virology, Chinese Academy of Sciences

WVU and Marshall University Combined Genomics Core Facilities

Wyoming Public Health Laborator

Wyoming Public Health Laboratory

Yale School of Public Health

Zhejiang Provincial Center for Disease Control and Prevention

Zoonotic and Exotic infection Diseases Division, Harbin Veterinary Research Institute, CAAS

Zoonotic and Exotic infection Diseases Division, Harbin Veterinary Resrarch Institute, CAAS

Zoontic and Exotic infection Diseases Division

Zooprofilattico Sperimentale dell'Emilia Romagna e della Lombardia (IZSLER), Risk Analysis and Genomic Epidemiology Unit

Zurita & Zurita Laboratorios
