## Supplementary material for "Preliminary report on SARS-CoV-2 Spike mutation T478K": SuppTable1_european.docx

| Country | S.T478K.Mutations | Total.Sequences | Percentage.over.total |
| --- | --- | --- | --- |
| Germany | 216 | 64531 | 0.33 |
| Switzerland | 170 | 29863 | 0.57 |
| Sweden | 162 | 23076 | 0.7 |
| United Kingdom | 116 | 380491 | 0.03 |
| France | 57 | 21573 | 0.26 |
| Netherlands | 50 | 23022 | 0.22 |
| Spain | 30 | 20017 | 0.15 |
| Turkey | 28 | 3561 | 0.79 |
| Belgium | 21 | 15604 | 0.13 |
| Bulgaria | 6 | 1423 | 0.42 |
| Denmark | 4 | 50604 | 0.01 |
| Italy | 4 | 21968 | 0.02 |
| Portugal | 4 | 5741 | 0.07 |
| Lithuania | 3 | 5963 | 0.05 |
| Luxembourg | 3 | 5943 | 0.05 |
| Austria | 1 | 4107 | 0.02 |
| Finland | 1 | 3045 | 0.03 |
| Kosovo | 1 | 37 | 2.7 |
| Poland | 1 | 6338 | 0.02 |
| Russia | 1 | 2880 | 0.03 |
