## Supplementary figures and images for "Preliminary report on SARS-CoV-2 Spike mutation T478K"

### SuppFigure1_lineages.png

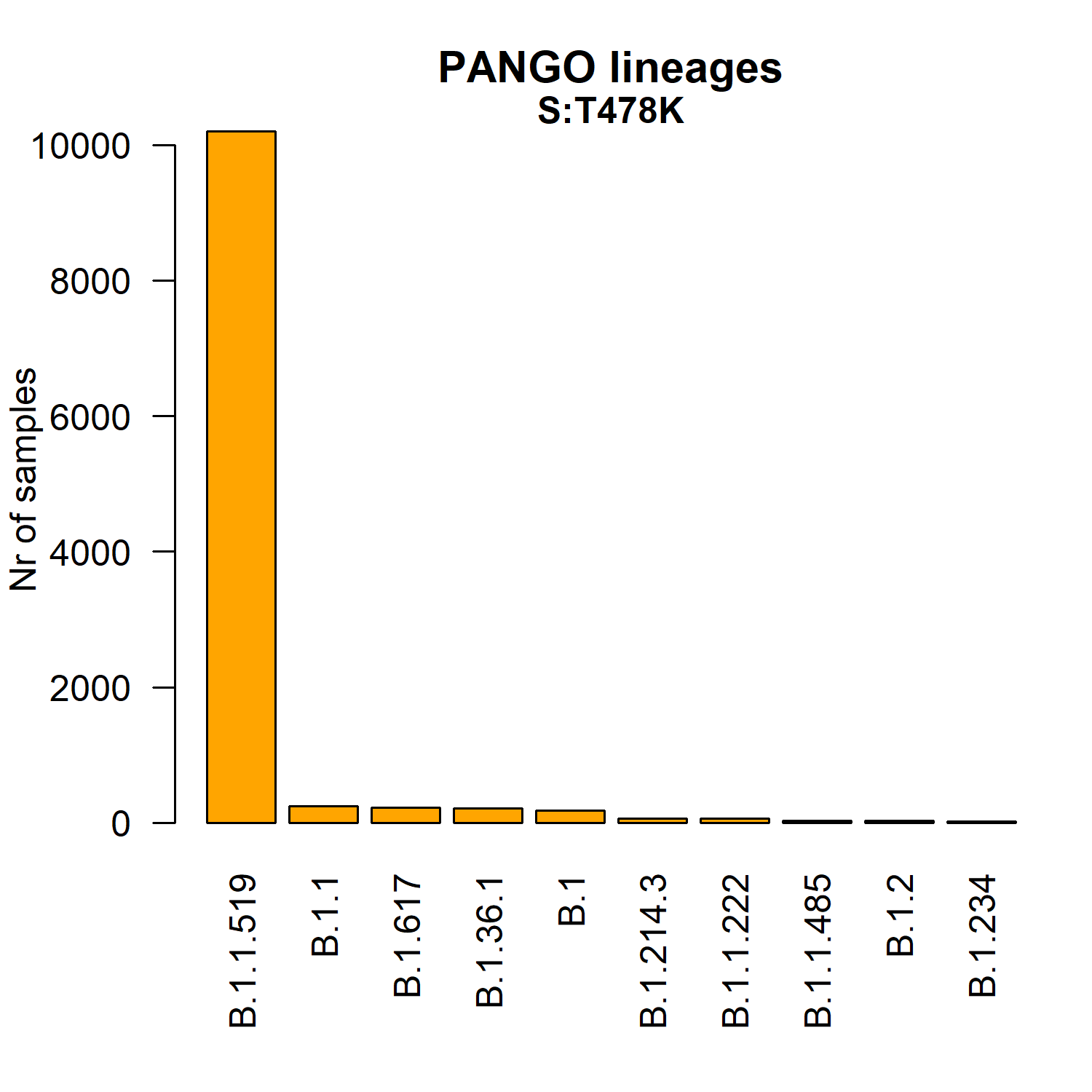

### SuppFigure2_age.png

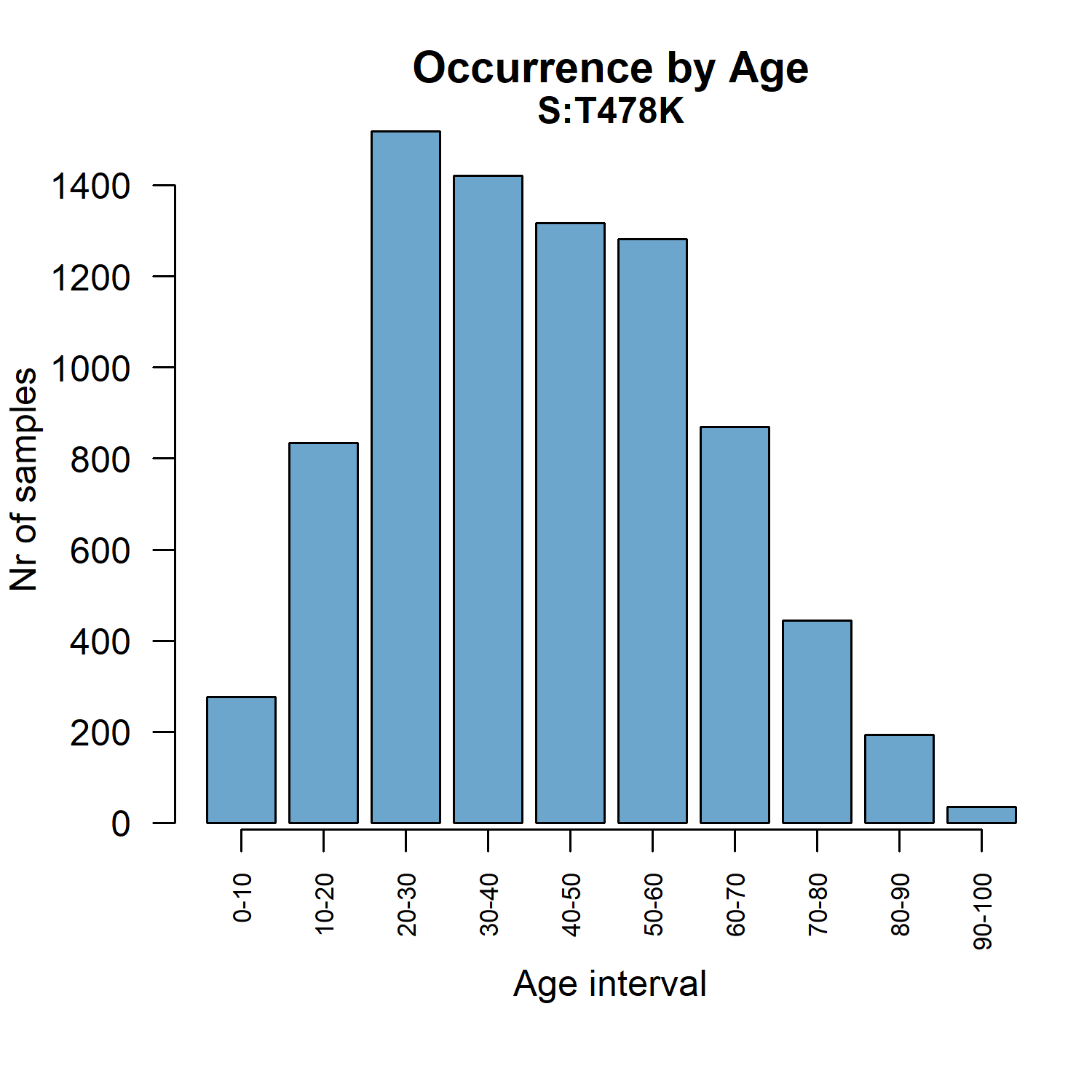

### SuppFigure3_countries.png

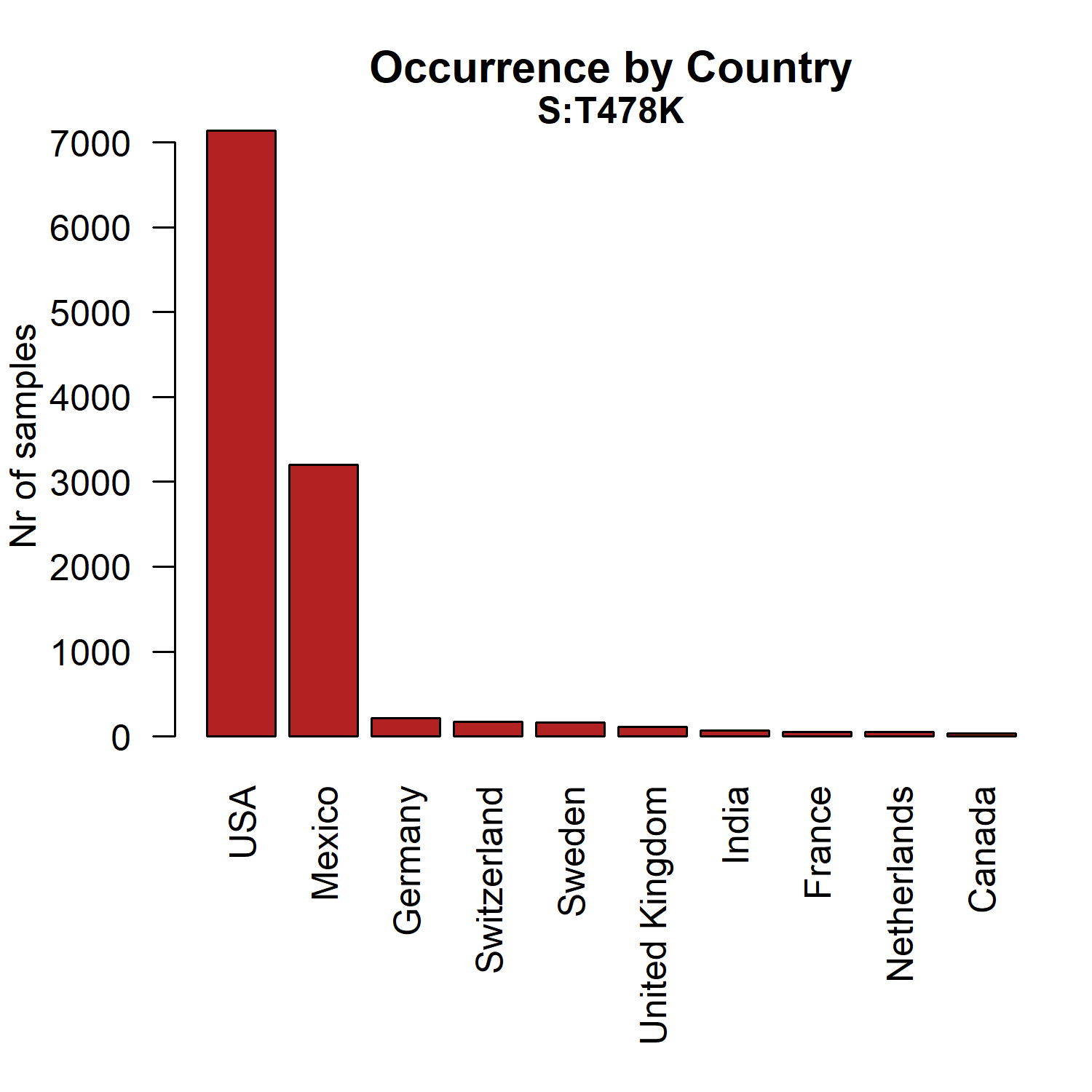

### SuppFigure4_world_frequency.png

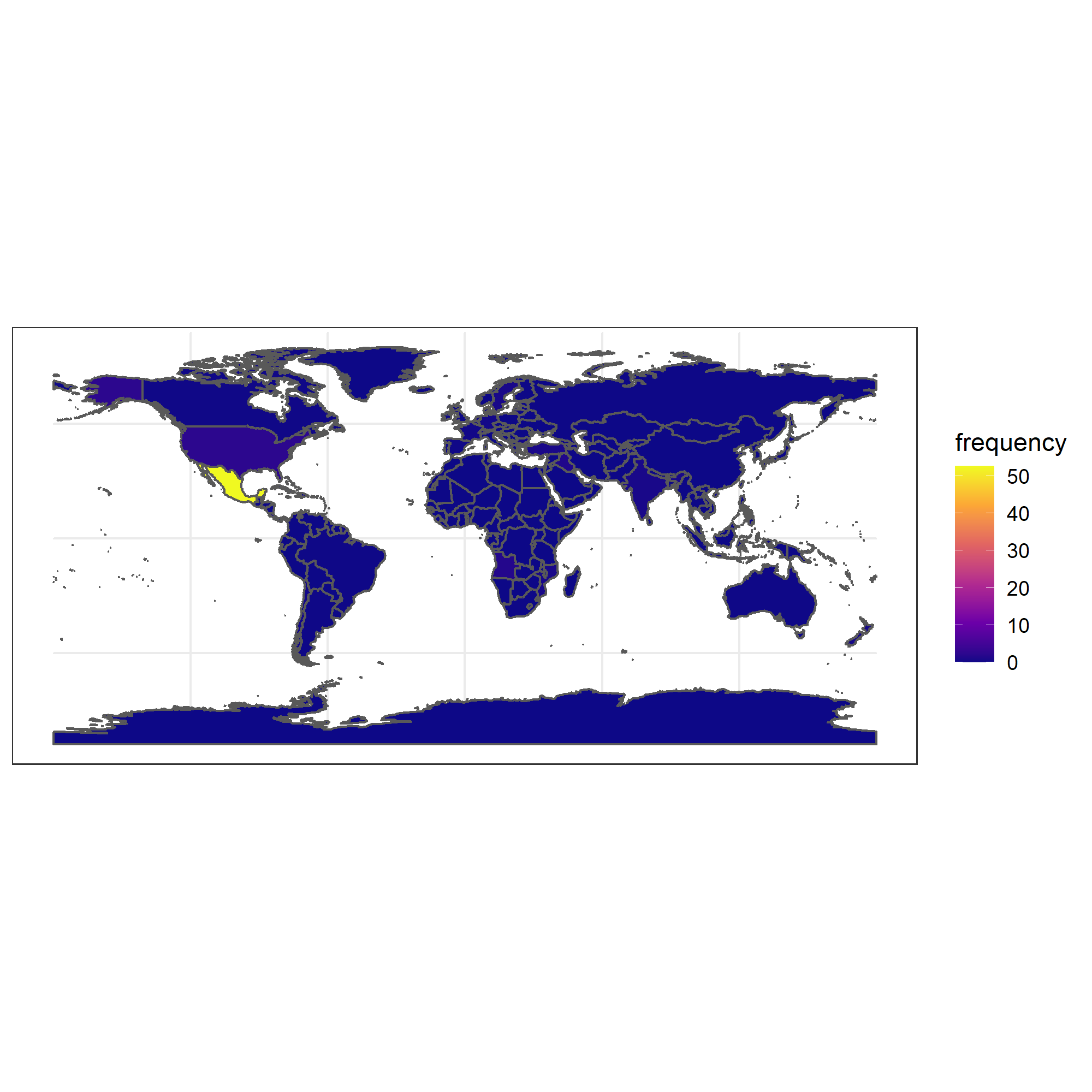
